# CaMKII oxidation is a performance/disease trade-off in vertebrate evolution

**DOI:** 10.1101/767525

**Authors:** Qinchuan Wang, Erick O. Hernández-Ochoa, Meera C. Viswanathan, Ian D. Blum, Jonathan M. Granger, Kevin R. Murphy, An-Chi Wei, Susan Aja, Naili Liu, Corina M. Antonescu, Liliana D. Florea, C. Conover Talbot, David Mohr, Kathryn R. Wagner, Sergi Regot, Richard M. Lovering, Mark N. Wu, Anthony Cammarato, Martin F. Schneider, Gabriel S. Bever, Mark E. Anderson

## Abstract

Reactive oxygen species (ROS) contribute to health and disease. CaMKII is a widely expressed enzyme whose activation by oxidation of regulatory domain methionines (ox-CaMKII) contributes to cardiovascular disease, asthma, and cancer. Here we integrate comparative genomic and experimental data to show that CaMKII activation by ROS arose more than half-a-billion years ago on the vertebrate stem lineage where it constituted a bridge between ROS and increased intracellular Ca^2+^ release, exercise responsive gene transcription, and improved performance in skeletal muscle. These enhancements to fight-or-flight physiology were likely key in facilitating a well-evidenced shift in the behavioural ecology of our immediate chordate ancestors, and, in turn, the evolutionary success of vertebrates. Still, the ox-CaMKII innovation for augmenting performance must be considered a critical evolutionary trade-off, as it rendered us more susceptible to common and often fatal diseases linked to excessive ROS.

---

The Ca^2+^-and calmodulin-dependent protein kinase II (CaMKII) is a multifunctional enzyme that augments intracellular Ca^2+^ flux, and regulates gene transcription^1^. CaMKII is initially activated by binding calmodulin, but post-translational modifications^2–5^ of conserved regulatory domain residues convert CaMKII into a Ca^2+^- and calmodulin-independent conformation by preventing the regulatory domain from occluding the active site (Fig. 1a). These post-translational modifications include autophosphorylation at threonine 287 (T287, numbered in accordance with CaMKIIγ isoform)^4,5^, *O*-GlcNAclyation at serine 280 (S280)^3^, and oxidation at cysteine 281/methionine 282 (CM; CaMKIIα) or methionine 281/282 (MM; CaMKIIγ, δ and β)^2^. CaMKII activity using the oxidative pathway (ox-CaMKII) is elevated in tissues of patients with cardiovascular diseases^6,7^, human cancer cell lines^8^, and asthmatic human pulmonary epithelium^9^, suggesting ox-CaMKII contributes to common human diseases. Mutant knock-in mice where the MM module in CaMKIIδ was replaced by ROS-resistant VV (valine 281/282) mutations are protected against a range of diseases that involve elevated oxidative stress, including stroke, cardiac arrhythmia, ischemia-reperfusion injury, sudden death, and asthma^6,7,10–13^. Given its clinical relevance, we sought to better understand the biological role of ox-CaMKII by exploring its evolutionary origins and functional implications.

**Fig. 1.**
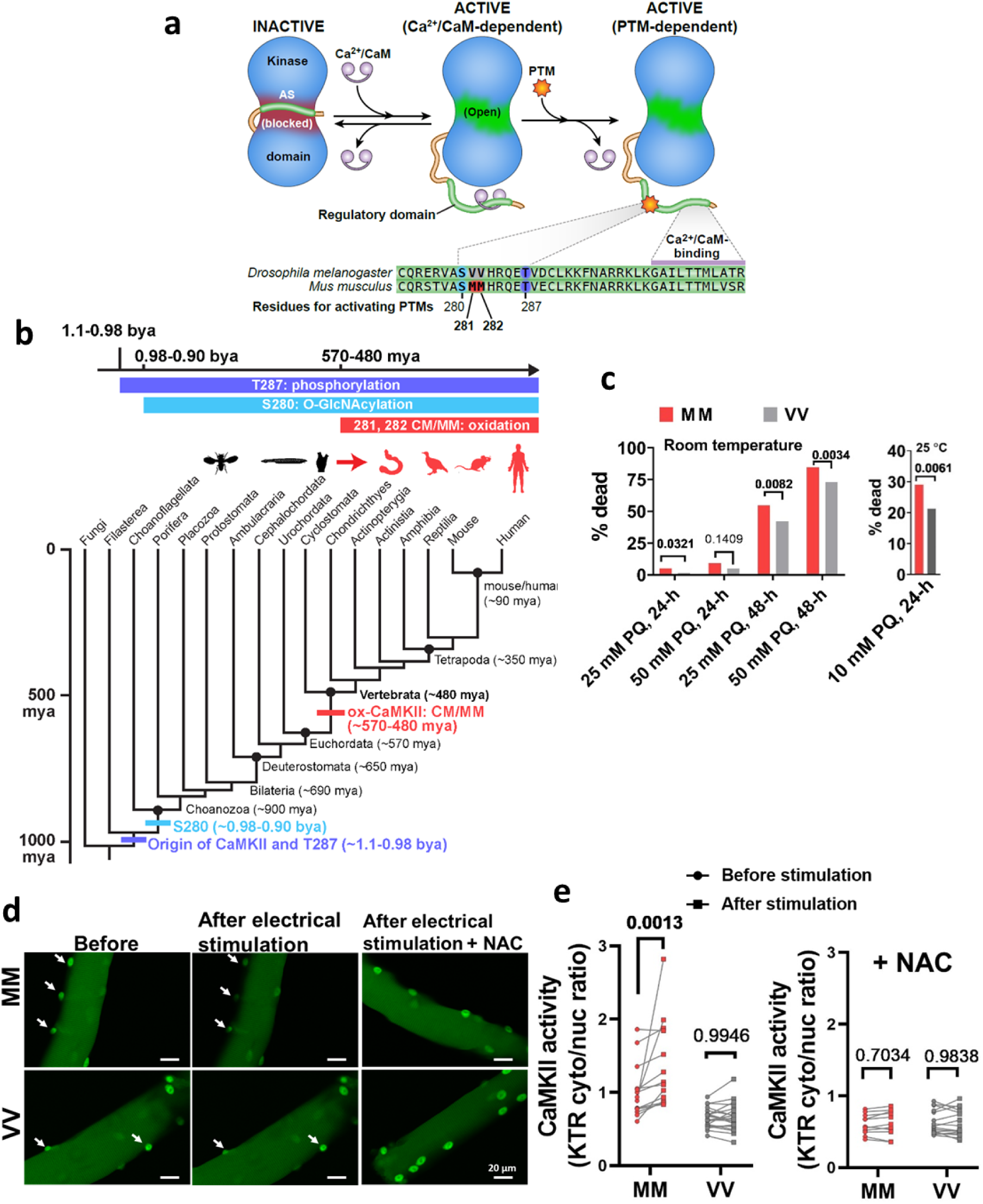
The MM motif in CaMKII arose in vertebrates, allowing CaMKII activation by ROS. **a**, CaMKII activation is initiated by binding Ca^2+^/CaM, but Ca^2+^/CaM independent CaMKII activity is sustained by post-translational (PTM) modifications of the regulatory domain. The CaMKII regulatory domain sequences of *Drosophila melanogaster* CaMKII and *Mus musculus* CaMKIIγ are shown. **b**, Phylogenetic origin and conservation of key residues for the activating PTMs of CaMKII (divergence time estimates from Kumar et al^50^). mya: million years ago; bya: billion years ago. **c**, Replacing the VV residues of *Drosophila melanogaster* CaMKII with MM increased mortality caused by exposure to 25 mM and 50 mM paraquat (PQ) incorporated in 5% sucrose solution at room temperature (21 °C, left panel) after 24 hours (24-h) and 48 hours (48-h), and by exposure to 10 mM paraquat at 25 °C after 24 hours (right panel). Only one out of 1020 flies fed control food (5% sucrose solution) died (not shown). *P*-values from Fisher’s exact test are shown above the square brackets. N = 216 and 217 respectively for MM and VV flies treated at the room temperature; n = 510 for both genotypes of flies treated by either control or paraquat solutions under the 25 °C test condition. **d**, Representative confocal micrographs of MM and VV mouse FDB muscle fibres expressing CaMKII-KTR before and after field electrical stimulation in the absence or presence of 2 mM N-acetylcysteine (NAC). Arrows indicate nuclei. **e**, Quantification of CaMKII activity (cytosolic to nuclear CaMKII-KTR signal ratio) in MM and VV fibres before and after field stimulation in the absence or presence of 2 mM NAC (Two-way ANOVA followed by Sidak’s multiple comparisons test. n = 14 nuclei from 6 fibres (MM) and n = 21 nuclei from 7 fibres (VV) in the left panel, and n = 11 nuclei from 5 fibres (MM) and n = 16 nuclei from 6 fibres (VV) in the right panel. *P*-values are shown above the brackets).

## MM residues confer ROS sensitivity to CaMKII in vertebrates and invertebrates

The phylogenetic distribution of ox-CaMKII supports the inference that an oxidation-sensitive amino acid pair in the regulatory domain is a derived feature of vertebrates, one that emerged on the vertebrate stem lineage sometime between approximately 570 and 480 million years ago (Fig 1b; Extended Data Fig. 1). This origin contrasts markedly with that of the T287 and S280 regulatory pathways. These appear to have evolved concurrent with CaMKII itself and are features ancestral to all metazoans, antedating the CM module by an additional ∼500 million years (Fig 1b). The consistent expression among crown vertebrates of an oxidation-sensitive pair at regulatory loci 281/282 indicates this vertebrate oxidative pathway has never been lost. It also indicates that ox-CaMKII is likely integrated within functionally and evolutionarily beneficial cascades, but at the cost of enhanced susceptibility to common, ROS-mediated diseases.

To test whether the MM module was sufficient to convert an invertebrate CaMKII into a ROS sensor, we mutated the ROS-resistant VV residues in *Drosophila melanogaster* into MM (Extended Data Fig. 2). *Drosophila melanogaster* is advantageous as a model, in part, because it has only one *CaMKII* gene. We found that the *CaMKII^MM^/CaMKII^MM^* flies (referred to as MM flies hereafter) showed a significantly higher mortality when fed sucrose solutions dosed with paraquat, a ROS-inducing toxin^14^ (Fig. 1c). These results establish a phylogenetically justified inference that the evolutionary appearance of these residues did confer ROS sensitivity through CaMKII activity in our stem-vertebrate ancestors.

The vertebrate stem lineage was witness to a major shift in behavioural ecology that set the stage for the modern (crown) vertebrate radiation and eventually our own evolutionary origin^15^. Out was the sessile, filter-feeding existence that served our deuterostome ancestors well and that still characterizes the adults of our closest living chordate relatives — the lancelets and tunicates. In was the metabolically costly strategy of being an active marine predator. A large series of structural innovations supported this dramatic transition by fundamentally altering the way in which these stem vertebrates received, processed, and acted on environmental stimuli. A few examples include: a prechordal head with complex organs of special sensation and a fully differentiated forebrain, a cartilaginous internal skeleton, muscular pharyngeal arches supporting advanced respiration, a more powerful heart pumping blood containing haemoglobin-rich red-blood cells, and a sympathetic nervous system and elaborate endocrine glands supporting a fight-or-flight physiological response. Many of these novel functional complexes were developmental products of a newly evolved population of highly migratory and multipotent neural crest cells, and, ultimately, the entire suite of derived vertebrate features may well owe its existence to a greatly expanded genetic tool kit made available by one, and possibly, two full rounds of genome duplication^16^. At some level, all of these innovations promote an increasing level of activity that must be enacted through the skeletal (striated) musculature; so, it was with the physiology of skeletal muscle that we tested for potential beneficial effects of ox-CaMKII.

CaMKII activity contributes to skeletal muscle function^17,18^, so we hypothesized that gaining the MM motif allowed ROS to enhance skeletal muscle performance through ox-CaMKII. We set out to test whether MM residues could dynamically respond to ROS in muscle fibres to increase CaMKII activity. To measure the dynamic change of CaMKII activity in muscle fibres, we developed a fluorescent reporter that translocates from the nucleus to the cytosol in response to increased activity of CaMKII (kinase translocation reporter, or KTR, Extended Data Fig. 3a and b and Methods)^19^. We validated the reporter in RPE-1 cells by showing that it translocated into the cytosol when the cells were treated by histamine (Extended Data Fig. 3c and d), which transiently increased cytosolic Ca^2+^ concentration^20^ (data not shown). This translocation was enhanced by co-expressed exogenous CaMKII, and blunted by co-expressed kinase-dead CaMKII mutant^21^ (CaMKII^K43M^) and by the CaMKII-specific inhibitory protein CaMKIIN^22^, supporting that the KTR translocation was driven by CaMKII activity. The results show that the cytosol/nuclear distribution of the KTR is a sensitive measurement of cellular CaMKII activity (Extended Data Fig. 3e). To determine the role of MM residues for ROS-induced CaMKII activity in muscle fibres, we developed a knock-in mouse where the MM residues of CaMKIIγ were replaced with VV (Extended Data Fig. 4a and 4b, homozygous *CaMKII*γ*^VV^/CaMKII*γ*^VV^* mice are referred to as VV mice hereafter). We selected *Camk2g* as the background because we found that CaMKIIγ is the most abundant isoform in mouse skeletal muscle (Extended Data Fig. 5a). The knock-in mutation did not change the mRNA expression from the *Camk2g* gene (Extended Data Fig. 5b). We introduced the CaMKII-KTR into the flexor digitorum brevis (FDB) skeletal muscles of MM (referring to wild type or WT mice) and VV mice by electroporating plasmids^23^ encoding the reporter and isolated the FDB muscle fibres after KTR was expressed (see Methods). We found that the increase of CaMKII activity in response to electrical stimulation required the MM motif (Fig. 1d and e). Addition of the antioxidant N-acetylcysteine (NAC) eliminated the rise in CaMKII activity in both VV and MM (WT) FDB fibres in response to stimulation (Fig. 1d and e). The results showed that ROS contribute to the activation of myofibre CaMKII, a process dependent on the MM module of CaMKII.

## Ox-CaMKII promotes exercise performance

Based on our hypothesis that ox-CaMKII arose in vertebrates to enhance skeletal muscle performance, we next asked whether loss of the MM residues affected exercise by testing the mice with maximal coerced treadmill exercise (Fig. 2a). MM (WT) mice ran farther (Fig. 2b), and attained higher maximal speeds (Fig. 2c) compared to VV littermates. Although oxidative stress and the MM motif were necessary for normal CaMKII activity in stimulated muscle fibres (Fig. 1d and e), the reduced exercise performance in VV mice could be due to muscle extrinsic factors, such as motivation or metabolism. Blood lactate accumulation correlates with perceived effort during progressively intensifying exercise^24^. However, we found no difference in blood lactate concentration between MM (WT) and VV mice either before or after running (Extended Data Fig. 6a), suggesting that perceived effort was similar. Furthermore, the VV and MM (WT) mice engaged equally in voluntary wheel running (Extended Data Fig. 6b), suggesting that the difference in forced running performance in VV mice was unlikely a consequence of reduced motivation for running. Exercise demands uninterrupted energy supply, and the depletion of blood glucose can be a limiting factor for endurance running in mice^25^. Furthermore, CaMKIIγ promotes hepatic gluconeogenesis^26^, an important source of blood glucose during exercise^27^. However, we found no difference in blood glucose between MM (WT) and VV mice before or immediately after running (Extended Data Fig. 6c), suggesting that the MM module does not contribute to blood glucose maintenance under these conditions. In addition, we found no significant difference in the body weight, lean mass, and fat mass between MM (WT) and VV mice; nor did we find a significant difference in their activity inside cages, food intake, oxygen consumption rate (VO_2_), CO_2_ production rate (VCO_2_), respiratory control ratio (RER), or energy expenditure (data not shown) when the mice were monitored individually in the Comprehensive Lab Animal Monitoring System (CLAMS), suggesting that the energy metabolism of MM (WT) and VV mice are similar. Based on these negative findings, we next determined whether the diminished exercise performance observed in VV mice might result from reduced muscle function *in vivo*. We determined the reduction in quadriceps contractility of repeated maximal isometric contractions elicited by direct repetitive electrical stimulation of femoral nerves, in anesthetized mice, as a measure of muscle fatigue (Fig. 2d). This *in vivo* approach allows for the direct examination of muscle function at body temperature, with intact blood flow and neuromuscular communication, while minimizing potential confounding factors derived from circulatory and nervous system feedback^28^. The VV mice exhibited earlier and enhanced fatigue compared to MM (WT) littermate mice (Fig. 2e and 2f). The reduced performance of the VV mice was unlikely due to developmental defects or gross pathological remodelling, as we found that the VV and MM (WT) mice have similar muscle weight to body weight ratios and grip strength (Extended Data Fig. 7a-g). In addition, the contents of mitochondrial complexes in muscles (Extended Data Fig. 8a), and oxidative phosphorylation and glycolysis capacities of isolated FDB muscle fibres were all similar between MM (WT) and VV mice (Extended Data Fig. 8b and c). Furthermore, we found no significant change in the fatigue-resistant type I fibres, and noted significant but subtle switches among type II fibres in VV quadriceps muscles, which were unlikely to explain the reduced endurance capacity (Extended Data Fig. 9a and b). Taken together, these data support a view that ox-CaMKII enhances dynamic responses to exercise and skeletal muscle performance.

**Fig. 2.**
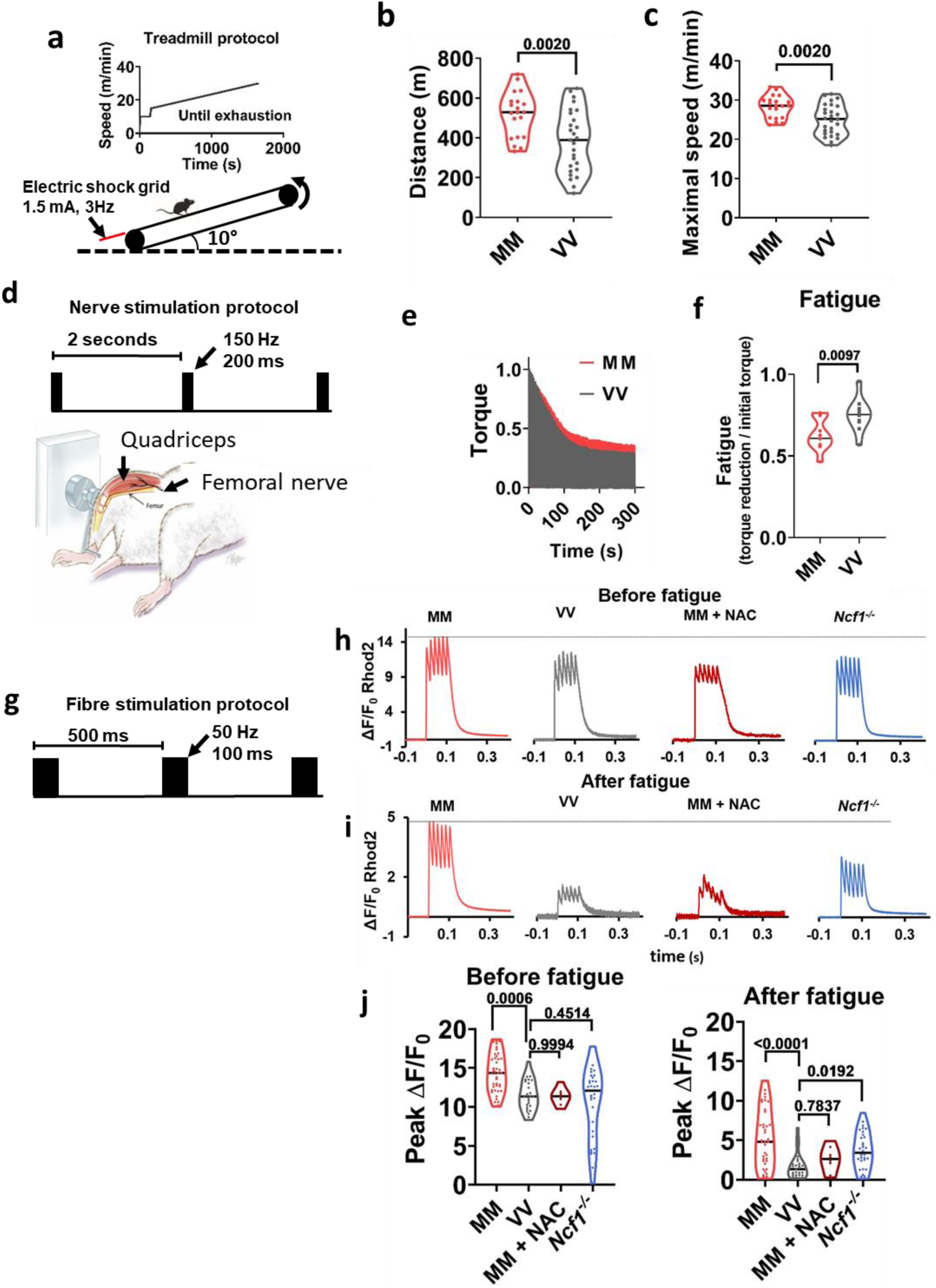
Ox-CaMKII supports exercise performance and enhances Ca^2+^ transients in mouse skeletal muscle fibres. **a**, Protocol for treadmill exercise. **b**, Running distance, and **c**, maximal speed attained prior to exhaustion. Unpaired Student’s *t* test, two-tailed, *P*-value shown above the square brackets, n = 20 MM (WT) mice and n = 27 VV mice. **d**, Experimental apparatus and electrical stimulation protocol for assessing quadriceps muscle performance *in vivo*. **e**, Averaged traces of quadriceps torque (normalized against maximum) from MM and VV mice during optimized nerve stimulation, n = 10 VV mice, and n = 11 MM mice. **f**, Quantification of fatigue defined by torque reduction divided by initial torque of each individual mouse, as shown in e. Unpaired Student’s *t* test, two-tailed, *P*-value shown above the square bracket. **g**, Protocol for field electrical stimulation of isolated FDB muscle fibres loaded with the Ca^2+^ sensitive fluorescent dye Rhod2. Rhod2 fluorescence during one cycle of electrical stimulation in MM and VV fibres, an MM fibre treated by NAC, and a *Ncf1^−/−^* fibre, before (h) and after (i) fatigue. **j**, Quantification of peak Ca^2+^ transients as measured in h and i. Before fatigue: n = 42 MM, n = 27 VV, n = 8 MM + NAC, and n = 32 *Ncf1-/-* fibres; after fatigue: n = 40 MM, n = 27 VV, n = 8 MM + NAC, and n = 32 *Ncf1-/-* fibres; *P*-values are shown above the brackets; Dunnett’s multiple comparisons tests comparing all other groups to VV fibres. Horizontal lines in b, c, f, and j indicate medians.

Intracellular Ca^2+^ grades myofilament interactions, thereby serving as an essential signal for muscle performance, and CaMKII promotes intracellular Ca^2+^ release in excitable tissues, including skeletal muscle^17,18^. We used a validated *in vitro* model of skeletal muscle fibre fatigue^29^, under conditions where we monitored the intracellular Ca^2+^ transients (see Methods, Fig. 2g). The VV fibres had reduced Ca^2+^ transients under basal conditions compared to MM (WT) counterparts (Fig. 2h). Fatigue is marked by reduced intracellular Ca^2+^ transients^30^, and VV fibres showed significantly greater reductions in these Ca^2+^ transients compared to MM (WT) (Fig. 2i). In order to test whether the exaggerated fatigue Ca^2+^ phenotype in VV muscle fibres was a consequence of ROS-signalling, we treated MM (WT) fibres with NAC. The MM fibres exposed to NAC phenocopied the Ca^2+^ release profiles measured in VV fibres (Fig. 2h-j). We next used a genetic approach to reduce ROS by isolating muscle fibres from *Ncf1*^−/−^ mice that lack p47^31^, an essential protein cofactor for NADPH oxidases that are an important source of ROS in skeletal muscles^32–34^. Similar to NAC treated MM (WT) fibres, the *Ncf1^−/−^* fibres shared a phenotype of diminished Ca^2+^ transients resembling VV muscle fibres (Fig 2. h-j). Taken together, we interpret these data as supporting a model where ox-CaMKII contributes to enhanced skeletal muscle performance, at least in part, by connecting ROS to mobilization of intracellular Ca^2+^.

## Ox-CaMKII regulates acute transcriptional responses to exercise

Exercise imposes metabolic, mechanical and redox stresses on skeletal muscle leading to transcriptional adaptation that is partly orchestrated by CaMKII^35,36^. We next measured transcriptional responses to submaximal exercise in skeletal muscles, comparing poly(A)^+^ transcriptomes by RNA sequencing from MM (WT) and VV littermate mice under identical conditions of speed, time, distance and feeding conditions (Fig. 3a, see Methods). Principal components analysis showed that sedentary MM (WT) and VV muscles had very similar transcriptional profiles (Fig. 3b). In contrast, transcriptional responses to exercise by VV muscles were present, but diminished compared to MM (WT) (Fig. 3b). We found that 582 genes were significantly up or down regulated (multiple-test false discovery rate-adjusted *q*-value < 0.05) in the MM (WT) samples, whereas only 216 genes reached the same threshold of *q* < 0.05 in the VV muscles (Fig. 3c and Supplementary table 1). Among the significantly changed genes in VV muscles, most (180, or 83%) were recapitulated by the MM (WT) muscles. To further compare the transcriptional responses of MM (WT) and VV muscles at the level of individual genes, we ranked the exercise-responsive genes identified in the MM (WT) muscles based on their log_2_ (fold change) values, as diagrammed (Fig. 3d, left panel), and plotted these genes in heat map palettes (Fig. 3d, middle panel). The changes of the same set of genes in the VV muscles were shown in separate palettes (Fig. 3d, right panel), but in the same order as that of the MM palettes. It is clear that the MM module had heterogeneous effects on the response of individual genes to exercise: most exercise responsive genes preserved their qualitative responses in VV muscles, while the up- or down-regulations of a small number of genes were completely blunted or even reversed. The results suggest that ox-CaMKII plays important, but specified, roles in the acute transcriptional response of skeletal muscles to exercise.

**Fig. 3.**
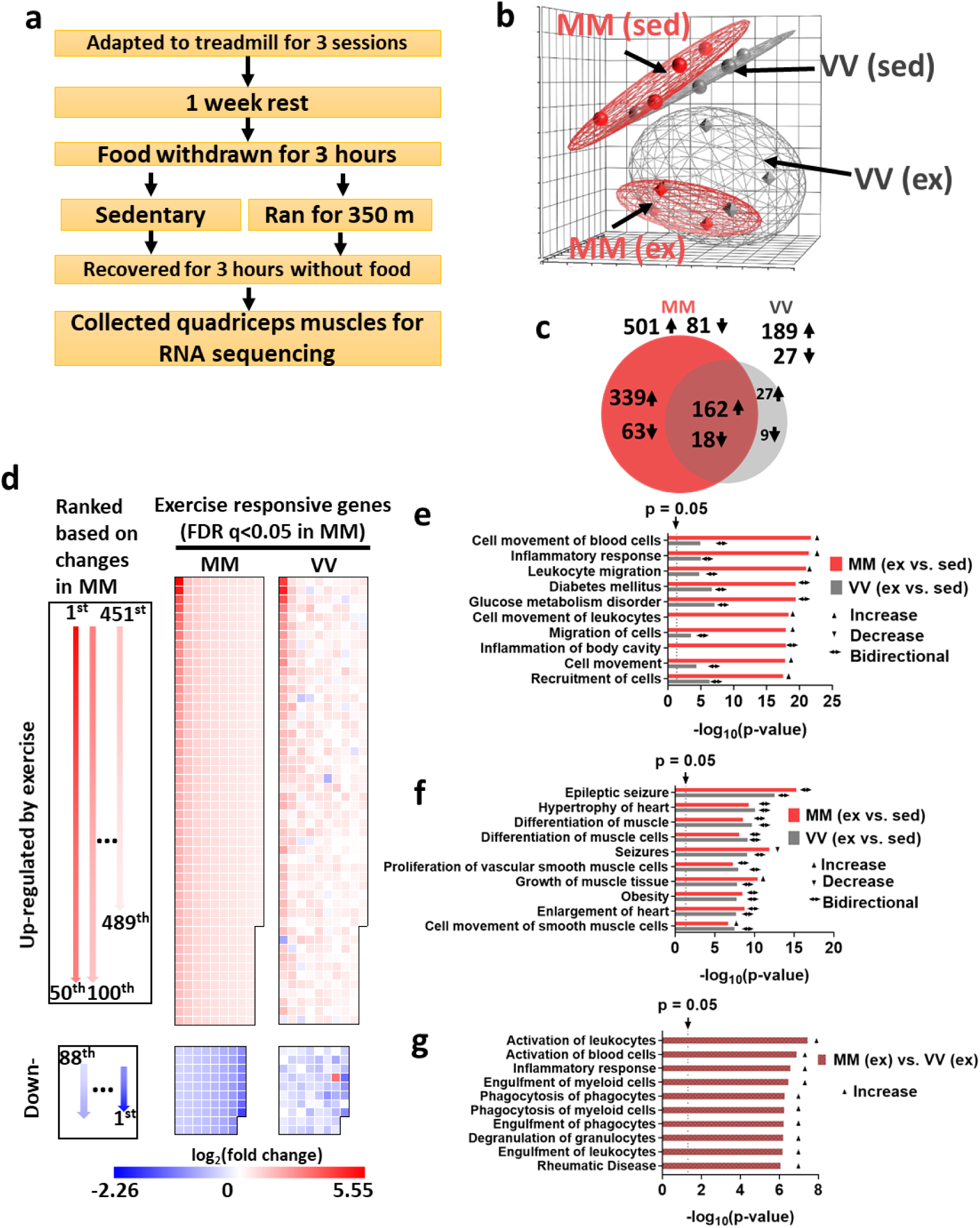
Ox-CaMKII is important for acute transcriptional responses to exercise. **a**, Protocol for submaximal exercise and muscle collection. **b**, Principal components analysis (PCA) of RNA sequencing results of sedentary (sed) and exercised (ex) muscle samples. The distance between samples in PCA corresponds to similarity (near) or difference (far) in their transcriptional profiles (n = 4 mice for each group). **c**, Numbers and overlap of significantly changed (false discovery rate-adjusted *q*-value < 0.05) genes in response to exercise in MM and VV muscles. Arrows indicates up- (↑) or down-regulation (↓) when comparing exercised muscles to sedentary muscles. Left panel of **d**, diagram of the layouts for arranging genes in the middle and right panels; middle panel of **d**, genes whose expression was significantly (*q* <0.05) changed in response to exercise in the MM (WT) muscles are ordered according to the diagram in the left panel, and their log_2_ (fold changes) are represented by colour; right panel, the log_2_ (fold change) of the same genes in response to exercise in the VV muscles are shown. **e**, Top-10 most significantly (smallest *P*-values) changed functions identified by Ingenuity Pathway Analysis comparing transcriptomes of exercised MM muscles to their sedentary counterparts. Corresponding enrichment *P*-values of the same functions in the exercised VV muscles are plotted for comparison. **f**, Top-10 most significantly (smallest *P*-values) changed functions identified by Ingenuity Pathway Analysis comparing transcriptomes of exercised VV muscles to their sedentary counterparts. Corresponding enrichment *P*-values of the same functions in the exercised MM muscles are plotted for comparison. **g**, Ingenuity pathway analysis directly comparing transcriptomes of exercised MM and VV muscles. In (e-g), activation, depression or bidirectional changes of the biological functions are determined by the z-score of Ingenuity Pathway Analysis for each pathway (z ≥ 2.0 for activation, z ≤ -2.0 for depression, otherwise for bidirectional changes).

We next used QIAGEN Ingenuity Pathway Analysis^37^ to extract biological pathway information from genes that showed a large shift (more than ±2σ) in expression in response to exercise. We found that in the MM (WT) muscles, eight out of the ten most significantly enriched biological function terms were related to inflammation (Fig. 3e). Strikingly, exercise induced lesser changes of the genes involved in the inflammatory response in the VV muscles (Fig. 3e). In contrast, the top 10 most enriched biological functions in VV muscles included terms such as “differentiation of muscle cells” and “growth of muscle tissue”, which were expected for the adaptive response to exercise^36^. Importantly, the MM (WT) muscles shared a similar pathway enrichment for these biological functions (Fig. 3f). We then directly compared the transcriptomes of exercised MM (WT) and VV muscles to identify a list of genes that showed the most prominent differences (more than ±2σ) between genotypes under this post-exercise condition. When these differentially regulated genes were analysed by Ingenuity Pathway Analysis, the results (Fig. 3g) further supported the prominent difference in inflammatory responses between exercised MM (WT) and VV muscles: exercised MM (WT) muscles showed significant enrichments (*p* < 0.05) and activation (z score ≥ 2.0) of multiple biological functions related to inflammation (Fig. 3g). Our results suggest that ox-CaMKII plays an important role in coupling ROS to the activation of physiological inflammatory response pathways, a well-established adaptive response to a single bout of unaccustomed exercise^38^. Under disease conditions CaMKII has been shown to promote inflammation in the heart^39–41^ and airway^11^, and to function in mast cells^11^, macrophages^42–45^ and T cells^46^. Our data unambiguously established ox-CaMKII as a molecular connection between inflammatory responses and physiological ROS signalling.

## MM enacted the performance/disease trade-off in flies

For it to be fixed by natural selection, the MM module likely provided fitness benefits to the ancestral vertebrates. Using *Drosophila melanogaster* as a model, we next tested whether the MM module could exert beneficial effects on physiological performance in an invertebrate. We reasoned that if a performance benefit is conferred by the MM module in flies it would suggest that the cellular context of invertebrates was permissive for the physiological benefits of ox-CaMKII. Since climbing involves insect leg muscles that are functionally and physiologically analogous to skeletal muscles of vertebrates^47^, we tested the climbing ability MM and VV (WT) flies. We placed the flies into vertical race-tracks too narrow for flying, but wide enough for climbing. When the flies were dislodged to the bottom of the race-tracks by vertical orientation of the apparatus, they climbed upwards as an innate escape response (Fig. 4a). Strikingly, the MM flies climbed at a significantly higher velocity than VV (WT) flies (Fig. 4b, control condition). The superior climbing performance conferred by the MM module was dependent on the physiological redox state, because ingesting food supplemented with the antioxidant NAC for 24 hours dose-dependently reduced the performance of MM but not VV (WT) flies (Fig. 4b, NAC-treated conditions). The MM flies climbed at similar velocities to VV (WT) flies after treatment by NAC (Fig. 4b). These results suggested that the MM module was capable of enhancing physiological responses to ROS in an invertebrate and likely also in ancestral vertebrates. To determine whether the MM module plays a direct role in *Drosophila melanogaster* striated muscles, we evaluated the performance of denervated hearts in MM and VV (WT) flies. In *Drosophila* the heart is a muscular tube that is experimentally accessible (Fig 4c). The hearts of MM flies had significantly better basal performance evidenced by significantly higher shortening velocity and relaxation rate (Fig. 4d and e). Similar to the vertical climbing assay, the performance benefits of the MM module in heart tubes were lost after exposure to NAC (Fig 4d and e). We interpreted these studies to show that ox-CaMKII was capable of producing ROS-driven physiological enhancements across evolutionarily distant species.

**Fig. 4.**
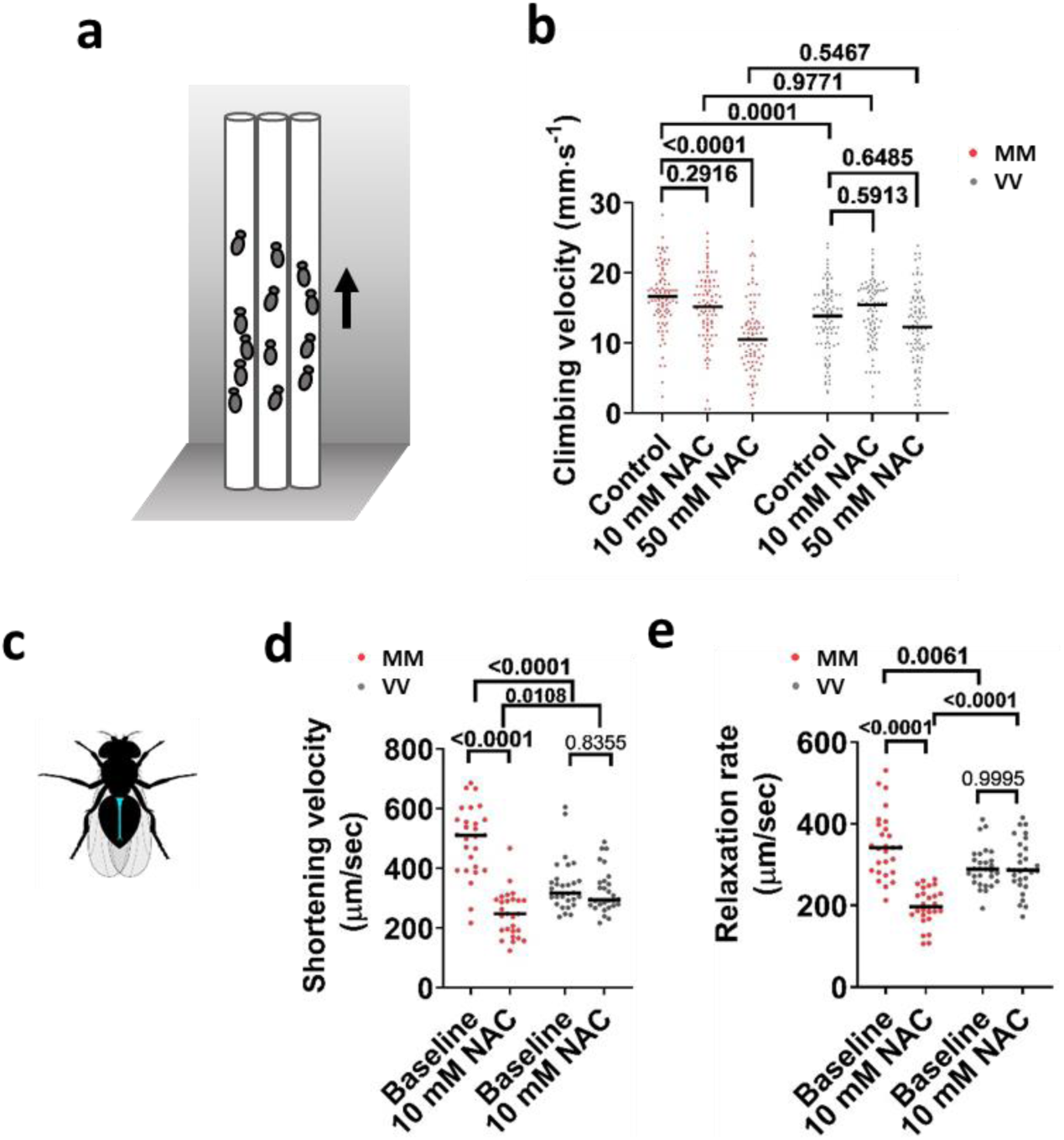
MM module couples ROS to improved performance in *Drosophila melanogaster*. **a**, Diagram for climbing test. **b**, Vertical climbing velocity of flies treated by control food (5% sucrose) or food containing 10 mM or 50 mM NAC for 24 hours; n = 93 control-treated MM, n = 90 10 mM NAC-treated MM, n = 86 50 mM NAC-treated MM, n = 94 control-treated VV, n = 91 10 mM NAC-treated VV, n = 94 50 mM NAC-treated VV. Horizontal lines indicate medians. *P*-values shown above the brackets, Tukey’s multiple comparisons test. **c**, diagram of a fly heart (in blue color). **d** and **e**, Cardiac performance indices of MM and VV hearts before and after 60-minute treatment by 10 mM NAC. The shortening velocity (d) and relaxation rate were assessed. n = 27 MM hearts per condition and n = 29 VV hearts per condition. Horizontal lines indicate medians. *P*-values shown about the brackets in (d) and (e), Tukey’s multiple comparisons test.

The striking benefits of MM modules on motor function and cardiac performance in flies (Fig. 4) contrasted with the fact that MM module promoted death when the flies were exposed to lethal doses of paraquat (Fig. 1c). Conceivably, when ROS increase above an optimal level, the MM module transduces the toxic ROS signal into excessive CaMKII activity that is detrimental to the flies. To test this possibility, we examined the effects of a sublethal dose of paraquat (4 mM for 24 hours) on climbing (Fig. 5a). After paraquat treatment, the MM flies exhibited significantly reduced climbing velocity, whereas VV (WT) flies were unaffected (Fig. 5a). The notion that excessive CaMKII activity is detrimental to motor function in invertebrates is supported by the observation that a hyperactive mutation of CaMKII impaired motor function of *Caenorhabditis elegans*^48^. We further examined the effects of very low dose paraquat feeding (1 mM for 3 to 6 days) on spontaneous ambulatory activity. We found that there was no difference in spontaneous ambulation between MM and VV (WT) flies at baseline, whereas exposure to food containing paraquat significantly reduced daily ambulatory activity counts only in MM flies (Fig 5b). Similarly, the benefits of the MM module on performance of denervated hearts were completely abrogated when the fly hearts were exposed to 10 mM paraquat for 90 minutes (Fig 5c and d); and longer exposure (150 minutes) to paraquat disrupted the contraction of significantly higher portions of MM than VV hearts (Fig 5e, and supplementary video 1). Taken together, effects of the MM module in fly CaMKII are strikingly similar to those in mice, strongly supporting the case for a performance/disease trade-off. The MM module promotes motor function in both species in response to physiological ROS, but switches from an asset to a liability when the organisms are challenged by pathological oxidative stress.

**Fig. 5.**
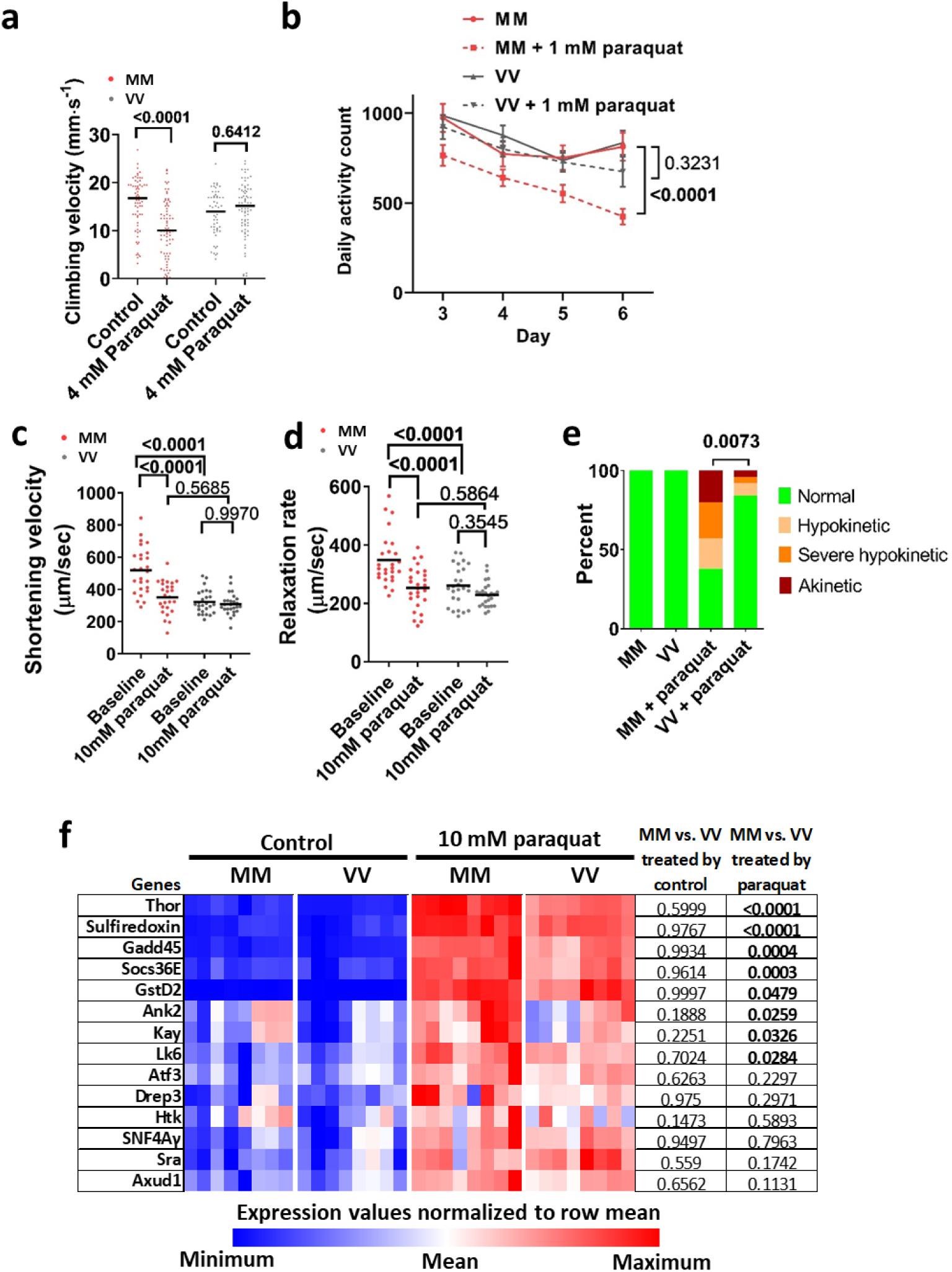
MM module sensitizes flies to pathological oxidative stress. **a**, Vertical climbing velocity of flies treated by control food or food containing 4 mM paraquat for 24 hours at 25 °C; n = 68 control-treated MM, n = 74 paraquat-treated MM, n = 57 control-treated VV, n = 75 paraquat-treated VV flies. Horizontal lines indicate medians. *P*-value from Sidak’s multiple comparisons test shown in the graph. **b**, Daily activity counts of MM and VV flies consuming a control or paraquat (1mM) diet. Diets started at day 1 and behaviour monitoring occurred between days 3 and 6 (Points and error bars are mean ± SEM. *P*-values calculated using Tukey’s multiple comparisons test for effect of paraquat, n = 29 control MM, n = 21 paraquat-treated MM, n = 31 control VV, and n = 21 paraquat-treated VV flies). **c and d,** Cardiac performance of hearts bathed first in control artificial haemolymph and then in haemolymph containing 10 mM of paraquat for 90 minutes. n = 26 hearts per genotype and the *P*-values from Tukey’s multiple comparisons test are shown. Horizontal lines indicate medians. **e**, All MM and VV hearts showed normal contraction before paraquat treatment, however, after exposure to 10 mM paraquat for 150 minutes, significantly more MM hearts became hypokinetic, severely hypokinetic or akinetic (examples of categorical cardiac performance are in the supplementary video 1). n = 26 hearts per genotype, *P*-value from Chi-square test. **f**, expression heat map of a subset of paraquat responsive genes that was quantified by RT-qPCR after RNA was extracted from flies ingesting control food (Control, 5% sucrose solution) or paraquat (10 mM of paraquat in 5% sucrose solution) for 24 hours at 25 °C (n = 8 biological replicates per group, each containing 15 males and 15 females). All of these genes were significantly upregulated by paraquat (*P* < 0.05, not shown, two-Way ANOVA). No genes showed significant differences between MM and VV flies fed control food, whereas a subset of genes had significantly higher expression in MM than in VV flies after exposure to paraquat (*P*-values shown in the table. Sidak’s multiple comparisons test).

Finally, we tested whether insertion of the MM module in CaMKII could establish unique connections between ROS and gene expression in flies, potentially mirroring the situation in mice (Fig. 3). Many mammalian transcription regulators targeted by CaMKII have orthologous counterparts in *Drosophila melanogaster* (http://flybase.org/). We fed our MM and VV (WT) flies with 10 mM paraquat for 24 hours at 25 °C, a regimen that induced elevated mortality in MM flies (Fig. 1c), and is known to induce a stereotyped transcriptional response in *Drosophila melanogaster*^49^. We focused on a subset of the paraquat-induced fly genes^49^ whose paralogues in mice were altered by exercise. We randomly selected some of these genes and confirmed by RT-qPCR that all were significantly regulated by paraquat in MM and VV flies (Fig. 5f). Strikingly, none of these genes showed a difference in expression between MM and VV flies consuming control food, while a subset exhibited significant differences between MM and VV flies after paraquat feeding (Fig. 5f). The results suggest that introducing the MM module to fly CaMKII bridges ROS to the expression of a specific set of genes, reminiscent of the transcriptional effects of the MM module in exercising mouse skeletal muscles (Fig. 3).

## Discussion

Our studies provide new *in vivo* and *in vitro* evidence that ox-CaMKII directly orchestrates connections between ROS, intracellular Ca^2+^ and gene transcription that lead to physiological advantages in mice, and, presumably, other vertebrates. Our pattern-based phylogenetic results indicate this advantage evolved concurrent with the establishment of the modern vertebrate body plan and its highly active behavioural ecology. It seems likely that ox-CaMKII was a key innovation in facilitating the heightened physiological output required of these derived anatomical systems and thus played a key role in the initial establishment and continued evolutionary success of vertebrates. The conservation of the CM/MM module in all isoforms of vertebrate CaMKII further suggests that ox-CaMKII plays diverse physiological roles, beyond those uncovered by this study in the skeletal muscles. The formative evolutionary role of the MM module is paired with considerable irony, given the well-recognized contributions of ox-CaMKII to major chronic and life-threatening human diseases. The striking observation that the MM module enacts the performance/disease trade-off in flies, an invertebrate diverged from our common ancestors for more than 500 million years is particularly worth noting. It suggests that the MM module is a concise but highly impactful ROS sensor, and once it was obtained by CaMKII in the ancestral vertebrate, the MM (CM) module was sufficient to couple ROS to a wide range of CaMKII targets important for enhanced performance, gene expression, disease and death. The totality of this information strongly supports the conclusion that the CM/MM module is an evolutionary trade-off in vertebrates that uses ROS to enhance physiological performance, while simultaneously bestowing sensitivity to ROS for promoting chronic diseases, many of which transpire in older organisms, beyond the reach of natural selection.

## Supporting information

Examples of normal and defective fly hearts

## Acknowledgement

We thank Drs. Hal Dietz and Gregg Semenza for their insightful comments and suggestions, Teresa Ruggle for assistance in graphic design, Benjamin Garlow for assistance in developing KTR, Jinying Yang for managing mice, and Tran Nguyen for maintaining fly stocks.

This work was supported by the National Institutes of Health (R35-HL140034 to MEA, R37-AR055099 to EOH and MFS, R01-AR059179 and R21-AR067872-01 to RML, R01-HL124091 to MCV and AC, R01-NS079584 to MNW) and a MOST grant (MOST-107-2636-B-002 -001 to MEA).

## Author contributions

Q.W., G.S.B. and M.E.A. contributed to the conception and design of the work; Q.W., E.O.H., M.C.V., I.D.B., J.M.G., K.R.M., A.W., S.A., N.L., D.M., K.R.W. and R.M.L. contributed to the acquisition of the data; Q.W., E.O.H., M.C.V., I.D.B., K.R.M., A.W., S.A., C.M.A., L.D.F., C.C.T., D.M., S.R., M.N.W., A.C. and M.S.F. analysed and interpreted the data; Q.W., G.S.B. and M.E.A drafted the manuscript; Q.W., G.S.B. and M.E.A. substantively revised the manuscript.

## Data availability statement

Raw sequencing data and the gene-expression matrix are available in the Gene Expression Omnibus (GEO) under accession number GSE132520. All other data are available from the corresponding author upon reasonable request.

## Methods

### Animal use

All animal handling procedures were in accordance with National Institutes of Health guidelines and were approved by the Institutional Animal Care and Use Committees of Johns Hopkins University School of Medicine.

### Generation of *CaMKII^MM^* point mutation in *Drosophila melanogaster* by CRISPR mediated gene editing

The genomic sequence of the *Drosophila melanogaster CaMKII* gene was used in the CRISPR guide design tool (http://crispr.mit.edu/) to create the CRISPR guides. The guide #1 (<u>GTTACAGCAACGCGAACGTG</u>) was chosen due to its close proximity to the codons encoding V281 and V282 and its lack of high probability off-targets. The guide #1 was ordered as complementary oligomeric DNA (Integrated DNA Technologies) and cloned into the pU6-BbsI-ChiRNA plasmid^46^ [the pU6-BbsI-chiRNA was a gift from Melissa Harrison & Kate O’Connor-Giles & Jill Wildonger (Addgene plasmid # 45946; http://n2t.net/addgene:45946; RRID:Addgene_45946)]. A single strand 176nt ultramer DNA oligo (ssODN-1R) was designed as the template for HDR-mediated point mutations and was ordered from Integrated DNA Technologies (Extended data Fig. 2a). The sequence of ssODN-1R is “AACATTGTCGTAAGTATGGCTCCCTTTAGCTTGCGCCGCGCATTAAATTTCTTGAGACAGTCTACGGTTTCTTGGCGATGCATCATGGAAGCGACTCGTTCGCGTTGCTGTAAC AATGTTTTTTCATTATCTTTATGTAAACCTAAGAGAAAAATTAGTCTGCACTTACACAAATC”. Injection of the ssODN-1R and the guide RNA encoding plasmids into fly embryos was carried out by Rainbow Transgenic Flies, Inc (3251 Corte Malpaso Unit 506 Camarillo CA). Genotyping was carried out by PCR amplification from genomic DNA, extracted from the wings of the flies, with primer-F (GTCGGTTATCCACCCTTTTG), and primer-R (GACGCCAAGTATATTGATGTGG) followed by Sanger sequencing and NsiI/NsiI-HF digestion. The flies with the correct *CaMKII^MM^* allele were backcrossed with iso31 flies^52^ for five generations to minimize the possibility of carrying off-targets from the CRISPR-mediated gene editing.

### Phylogenetic survey of CaMKII

Most CaMKII orthologues listed in Supplementary Fig. 1 were identified in the Interpro database (http://www.ebi.ac.uk/interpro/entry/IPR013543/taxonomy), based on the criteria that the sequences have the conserved CaMKII association domain, and a kinase domain. Additional sequences were uncovered by BLAST in the NCBL nucleotide database and translated into proteins (https://blast.ncbi.nlm.nih.gov/Blast.cgi). The CaMKII sequences were aligned using the Molecular Evolutionary Genetics Analysis software MEGA-X^53^ (https://www.megasoftware.net/). The evolution of an oxidation-sensitive amino acid pair at loci 281/282 of the CaMKII regulatory domain is an unambiguous synapomorphy of crown-clade vertebrates among Deuterostomata. The initial identity of this pair was CM, with the 281 cysteine likely being subsequently replaced by a methionine in one of the two paralogs that resulted from a full round of genome duplication that occurred prior to the origin of the vertebrate crown clade. Our phylogenetic survey did recover a small number of non-deuterostome taxa that also exhibit oxidizable residues at those same regulatory loci. The large phylogenetic separation between these taxa, both individually and collectively, from Deuterostomata leaves it clear that they evolved independent of the vertebrate condition; they are also MM rather than the CM of the earliest vertebrates. As a greater taxonomic diversity of metazoan genomes become available, a meaningful probabilistic analysis of ox-CaMKII evolution outside of Deuterostomata will be possible. But that analysis is highly unlikely to question the evolutionarily unique nature of vertebrate ox-CaMKII.

### Generation of *CaMKII^VV^* knock-in mutation in mice

*CaMKII^VV^* knock-in mice were generated by GenOway (https://www.genoway.com/) with mouse embryonic stem cells of the C57BL/6 background as specified in Extended Data Fig. 4. The mice used in experiments had been further backcrossed to C57BL/6J mice (The Jackson Laboratory, 000664) and all experiments were carried out with littermates.

### Western blotting for mitochondrial complexes

Protein extracts were prepared from frozen tissue with T-PER Tissue Protein Extraction Reagent (Thermo Scientific, #78510) in the presence of protease (Sigma-Aldrich, P8340) and phosphatase (Sigma-Aldrich, P0044) inhibitors. The primary antibodies were the Total OXPHOS Rodent WB Antibody Cocktail from Abcam (ab110413), and GAPDH (D16H11) XP® Rabbit mAb (Cell Signalling, #5174). Data were collected by LI-COR Odyssey Fc (Lincoln, Nebraska 68504 USA).

### Skeletal muscle fibre typing

Skeletal muscle fibre composition was determined by immunostaining following a standard method^54^. The primary antibodies BA-F8-c (myosin heavy chain, slow), SC-71-c (Myosin Heavy Chain Type IIA), BF-F3-c (Myosin Heavy Chain Type IIB), 6H1-s (myosin heavy chain, fast, IIX) were obtained from Developmental Studies Hybridoma Bank (University of Iowa).

### Paraquat treatment and behaviour study of flies

To determine the effects of paraquat on mortality, newly eclosed flies were sorted under CO_2_ anesthetization and placed into individual vials, and each vial received 10 males and 10 females. When the flies reached 5 to 7 days old, they were transferred into vials containing a filter paper pad (cut from Bio-Rad #1704085) soaked with 600 μL of 5% sucrose solution (control) or 5% sucrose solutions containing 10 mM, 25 mM or 50 mM paraquat (Sigma-Aldrich, # 856177). Mortality of flies was recorded at 24 and 48 hours after initiation of the treatment.

To test the negative geotaxis (climbing), females were collected and aged, as above, and kept in vials in groups of 10 flies. For low dose paraquat treatment, the flies were treated by 5% sucrose or 5% sucrose + 4 mM paraquat for 24 hours at 25 °C. They were then transferred into vertical test tracks made from 25 mL serological pipette tubes. During the climbing test, the flies were dislodged to the bottom of the tubes by rapidly tapping the vials on the desktop for 10 times and climbing was video recorded for subsequent analysis. Each group of flies was tested for 10 consecutive trials at 30 seconds intervals. The vertical distances the flies climbed in 6 seconds since the last tap (time 0) were used to calculate the vertical velocity of climbing. Flies that initiated flight or paused during the 6-second time window were excluded from the analysis. We found that the flies performed reproducibly from the second to tenth trials and presented data from trial 2 in Fig. 4b and Fig. 5a. To determine the effects of very low dose paraquat (1 mM) on daily ambulatory activity, individual 1-week old female flies were anesthetized by CO_2_ and loaded into tubes containing control or paraquat-containing food and monitored by *Drosophila* Activity Monitoring System (Trikinetics). Fly behaviour was recorded from day 3 to day 6.

### *Drosophila* cardiac physiological analysis

Dorsal cardiac tubes of ten-day-old female *CaMKII^WT^* (denoted VV (WT)) and *CaMKII^MM^* (denoted MM) flies were dissected in oxygenated artificial hemolymph^55^. Myogenic contractions of cardiac tissue were recorded using the Hamamatsu Orca Flash 2.8 CMOS camera on a Leica DM5000B TL microscope with a 10x immersion lens at ∼120 frames per second at baseline and after 90 and 150 minutes following the addition of 10 mM paraquat, or 60 minutes after the addition of 10 mM NAC. Cardiac physiological indices were determined using the semi-automated optical heartbeat analysis program^56,57^. Significant differences between genotypes before and after paraquat treatment for 90 minutes were determined by two-tailed Mann-Whitney tests. After 150 minutes in paraquat, many of the MM hearts no longer contracted and the cardiac indices could not be meaningfully derived. We, therefore, categorized the contractions as normal, hypokinetic (part of the heart contracting), severe hypokinetic (only twitching could be observed in part of the heart), and akinetic (no movement). Representative videos for each category are shown in supplementary video 1. The categorical data were assessed using a Chi-square test.

### Construction and validation of a CaMKII activity sensor CaMKII-KTR

The CaMKII-KTR was constructed based on the principles previously published^19^ and described in the Extended Data Fig. 3. Specifically, the sensor consists (from N-terminus to C-terminus) of a CaMKII-binding region derived from HDAC4, a linker, a nuclear localization signal (NLS), a nuclear exporting signal (NES) and a fluorescent protein. Optimized CaMKII phosphorylation sites were built into the NLS and NES while keeping the NLS and NES functional. The protein sequence of the CaMKII-KTR, excluding the enhanced green fluorescent protein, is EQELLFRQQALLLEQQRIHQLRNYQASMEAAGIPVSFGSHRPLKRTASVNEDEAPSKKPLARTASVSSRLERLTLQSS. A cDNA encoding this sequence was ordered as a codon-optimized gene block (gBlock_HDAC4-NLS-NES) from Integrated DNA Technologies (gccaccatgGAACAGGAACTGCTCTTCCGGCAACAGGCACTTCTGTTGGAGCAGCAACGAATCCATCAACTTAGAAACTACCAAGCATCAATGGAAGCAGCCGGGATTCCTGTCTCCTTCGGATCTCACAGACCTCTCAAAAGGACAGCTAGTGTAAACGAGGACGAAGCACCTTCAAAGAAACCCTTGGCTAGGACCGCTAGTGTCAGTAGTCGACTGGAGCGGTTGACACTTCAAAGTTCC). The gBlock_HDAC4-NLS-NES was cloned into Cerulean-N1 vector^58^ (Cerulean-N1 was a gift from Michael Davidson & Dave Piston, Addgene plasmid # 54742; http://n2t.net/addgene:54742; RRID:Addgene_54742) by In-fusion cloning technology (In-Fusion® HD Cloning Plus CE, Takara, CA). The Cerulean encoding region was then replaced by a stretch of cDNA encoding eGFP, derived from pEGFP-C1 (Takara, CA). To validate the response of the CaMKII-KTR to intracellular activity of CaMKII, we transfected (FuGENE® HD Transfection Reagent, Promega, WI) the CaMKII-KTR or co-transfected it with CaMKII, CaMKII^K43M^, and CaMKIIN constructs into RPE-1 cells and stimulated the cells with 50 μM histamine. Before cells were imaged, we replaced the culture medium with Live Cell Imaging Solution supplemented with 4.5 g/L glucose (ThermoFisher Scientific, A14291DJ), and stained their nuclei with Hoechst 33342 (ThermoFisher Scientific, #62249) for 20 minutes to facilitate identification of the nuclei. Fluorescent images were collected using an Olympus IX83 epifluorescence microscope equipped with an ORCA Flash 4.0 sCMOS camera and UPLSAPO20X NA0.75 objective lens. Cells were maintained at 37°C in an OkoLabs stage top incubator. Image analyses were carried out in CellProfiler^59^, which identified the nuclei and five-pixel-wide cytosolic rings surrounding the nuclei. The cytosolic to nuclear KTR signal ratios were calculated using the median intensities measured from the nuclei and cytosolic rings of individual cells.

### Mouse treadmill exercise

Exercise capacity tests were carried out with the Exer 3/6 Rodent treadmill (Columbus Instrument, Columbus, OH). Prior to exercise capacity testing, the mice (12 to 15-week-old) were acclimated to the treadmill for three sessions on three consecutive days. The treadmill was set to 10° inclination and the speed was set to 0, 5, and 10 m/min for the first, second and third acclimation sessions respectively. The electric shock grid at the rear end of the treadmill was turned on and set at stimulation intensity of 9 and frequency of 3 Hz. During exercise capacity testing, each mouse was placed into a lane of the treadmill. The genotype of the animals was blinded to the operator. The exercise protocol consisted of the following steps: (1) 10 m/min for 2 minutes for warm up, (2) continuous acceleration from 15 m/min at a rate of 0.6 m/min^2^ until the mouse was exhausted. Exhaustion was determined when the mouse stayed on the shock grid continuously for 5 seconds and was determined by the same observer for all experiments. Glucose and lactate were measured from a drop of blood from the tail tip before and immediately after exercise, using an OneTouch Ultra 2 glucometer (Lifescan, Inc) and a Lactate Plus lactate meter (Nova Biomedical) respectively.

### Mouse voluntary wheel running, and accompanying metabolic data

Voluntary wheel running data were collected from mice tested for 6 days in an open-circuit indirect calorimeter outfitted with running wheels (Comprehensive Lab Animal Monitoring System, Columbus Instruments) at the Center for Metabolism and Obesity Research service core. Data were collected continuously (Oxymax software, v.5.9, Columbus Instruments). Days 1-5 of acclimation to wheel running were monitored for expected daily increases in number of wheel rotations; the analysis of day 6 is presented. The instrument also provided data for voluntary physical activity in the main cage as indexed by counts of infrared beam breaks, intakes of powdered diet (2018, Envigo), as well as rates of O_2_ consumption (VO_2_, ml/kg/hr) and CO_2_ production (VCO_2_, ml/kg/hr). Oxymax software calculated the respiratory exchange ratio (RER = VCO_2_/VO_2_) to assess the oxidized fuel mixture being oxidized, and the rates of energy expenditure (EE, kcal/kg/hr; EE= VO_2_× [3.815 + (1.232 × RER)]. The standard outputs of indirect calorimetry data as per-kg/hr were also renormalized and analysed as per-kg-lean-mass/hr. Body composition data for lean mass and fat mass were obtained using an EchoMRI-100 at the Johns Hopkins University Phenotyping service core. None of the measures from these experiments showed group differences; data from voluntary wheel running are presented.

### Assessments muscle function *in vivo*

Grip strength measurements were carried out as described previously using a grip strength meter (Columbus Instruments, Columbus, OH, USA)^60^. Each mouse performed grip strength test until 6 successful attempts were accumulated, and the maximal force among the six attempts was taken as the grip strength.

*In vivo* quadriceps torque measurement was described previously^28^. Briefly, the mice were anesthetized under 4% isoflurane and then maintained at 1%. Then their pelvis, torso and femur were stabilized on the apparatus. Afterwards, the distal leg was taped to a lever arm, which was connected with a torque cell. The femoral nerve was stimulated subcutaneously to induce maximal quadriceps muscle contractions and the torque produced was recorded by a connected computer for subsequent analysis. The voltage of the stimulation was optimized prior to the studies to produce the maximal torque.

### *In cellulo* study of skeletal muscle fibres

Electroporation of DNA into flexor digitorum brevis (FDB) skeletal muscles of mice, muscle fibre culture, measurements of cytosol/nucleus distribution of CaMKII-KTR, and action potential-induced Ca^2+^ transient imaging followed our previous reports^23,29^. For N-acety-L-cysteine (NAC) treatment, the fibres were incubated for 20 minutes with 2 mM NAC (Sigma-Aldrich, St. Louis, MO; catalogue # A-7250). Fibres for study were randomly chosen, and where noted, 2-3 nuclei from the same fibre were studied.

For Seahorse study, FDB skeletal muscle fibers were isolated one day before the experiments and plated to a laminin-pretreated Seahorse XF96 Cell Culture Microplates overnight. Cell metabolism and bioenergetic analyses of muscle fibers were performed using an Agilent XF96 Extracellular Flux Analyzer. XF Cell Mitochondrial Stress Test kit and Glycolysis Stress Test kit were used to measure mitochondria respiration capacities and cellular glycolysis capacities following the manufacturer’s protocol. In the mitochondrial stress assay, muscle fibers were incubated in the muscle fiber assay medium (120 mM NaCl, 3.5 mM KCl, 1.3 mM CaCl_2_, 0.4 mM KH_2_PO_4_, 1 mM MgCl_2_, 5 mM HEPES, and 10 mM glucose, pH 7.4) followed by port injections of final concentration of 1 μM oligomycin, 0.5 μM FCCP, 10 mM pyruvate and 0.5 μM antimycin A/rotenone. In the glycolysis stress assay, glucose was not included in the initial muscle fiber assay medium, and then 2 mM glutamine, 10mM glucose, 1μM oligomycin, and 50mM 2-DG were injected sequentially. The oxygen consumption rate (OCR) and extracellular acidification rate (ECAR) were analyzed using Seahorse Wave software. OCR and ECAR were normalized by the number of skeletal muscle fibers per well.

### Mouse treatment and sample collection for RNA sequencing

Male mice between 13 and 15 weeks of age were used for the RNA sequencing experiments. The mice were first acclimated to the treadmill (Exer 3/6, Columbus Instrument, Columbus, OH) for 10 minutes on three consecutive days. The treadmill inclined at 10° and the speed was 0, 5 m·min^−1^, and 10 m·min^−1^ for day 1, 2 and 3 respectively. To minimize the effects of training on the skeletal muscles, the mice rested for 7 days before sample collection. On the day of sample collection, food was withdrawn from the mice at 9:00 AM to minimize the effects of food intake on signalling and gene transcription in the muscles. At 12:00 PM, the mice ran on a treadmill set to 10° of inclination. The treadmill speed was set at 10 m/min for 2 minutes to allow the mice to warm up. The speed was then increased to 15 m/min and then continuously ramped up from 15 m/min to 23 m/min at a rate of acceleration of 0.6 m·min^−2^. When the running protocol ended, the mice had run 350 meters, which was lower than the average running capacity of VV mice tested by the same treadmill protocol. Any mice that did not finish the protocol were excluded from subsequent sample collection. After exercise, the mice were allowed to rest for 3 hours with access to water but not food. Then they were euthanized by cervical dislocation after being anesthetized by isoflurane. The quadriceps muscles were quickly excised, frozen, and stored in liquid nitrogen.

### RNA extraction, quantification and quality control

To extract high quality total RNA, the quadriceps muscles were processed first in the Trizol reagent (ThermoFisher Scientific, Catalog # 15596018) and then purified by RNeasy mini columns (Qiagen, catalog # 74104) as follows. To avoid sampling bias, the entire quadriceps muscles were homogenized in Trizol reagent at the weight (mg) to volume (µL) ratio of 1:15. One mL of homogenate was processed following the manufacture’s protocol until the step of phase separation. Then, 0.5 mL of the aqueous phase was mixed with an equal volume of 70% ethanol for subsequent RNA purification by RNeasy mini kit (Qiagen, catalog # 74104) with on-column DNAse (RNase-Free DNase Set, Qiagen catalog # 79254) treatment. The concentration of the RNA was determined by Qubit fluorometric quantitation (Qubit RNA BR Assay Kit, ThermoFisher Scientific, catalog # Q10210). The integrity of the total RNA was determined by a Fragment Analyzer (Advanced Analytical Technologies, Inc). The average RNA Quality Number (RQN) was 9.04 ± 0.08 (Mean ± SEM, n=24).

### RNA sequencing library preparation

1 µg of total RNA from each sample was used for RNA sequencing library preparation with the TruSeq® Stranded mRNA Library Prep kit (Illumina, catalog # 20020594). The libraries were barcoded by TruSeq® RNA Unique Dual Indexes (Illumina, catalog # 20022371) and quantified by qPCR on a Bio-Rad CFX Connect Real-Time PCR detection system with the NEBNext Library Quant Kit for Illumina (New England Biolabs, catalog # E7630L). The libraries were normalized to 10 nM, pooled and sequenced.

### RNA sequencing and data analyses

RNA sequencing was carried out at Johns Hopkins School of Medicine Genetic Resources Core Facility on a NovaSeq 6000 sequencing system (Illumina) with a S1 flow cell for 200 cycles, generating 100 bp paired-end reads. Sequencing data processing was carried out by the Johns Hopkins Computational Biology Consulting Core. The statistics for sequencing data analysis are presented in supplementary tables 1-3. The RNA sequencing data were submitted to GEO repository and were assigned record number GSE132520.

Biological interpretation of sequencing results was carried out with the Ingenuity Pathway Analysis (IPA) platform (QIAGEN Ingenuity Systems, Redwood CA, USA) at the Johns Hopkins Deep Sequencing & Microarray Core Facility. The differentially expressed genes (over 2σ) between exercised and sedentary groups within the same genotypes and between the MM and VV samples after exercise were mapped by IPA to known pathways and biological functions of a curated Knowledge Base (QIAGEN Bioinformatics IPA Winter Release 2018). IPA evaluated the statistical significance of over-representation of the differentially expressed genes in each pathway or biological function, and a *P*-value < 0.05 was defined as statistically significant. In addition, IPA calculated the whether a pathway or biological function can be considered activated (z ≥ 2.0), inactivated (z ≤ -2.0) or bidirectional altered (-2.0 < z < 2.0) based the up- and down-regulation of genes involved the pathway or biological function.

### RT-qPCR

For samples from mice, we converted 1 μg of total RNA from each sample into cDNA with the iScript™ Reverse Transcription Supermix (Bio-Rad, Hercules, CA catalog # 1708840). 2 ng of cDNA were used in each qPCR reaction on a CFX Connect Real-time PCR detection system (Bio-Rad, Hercules, CA) with SsoAdvanced™ Universal SYBR® Green Supermix (Bio-Rad, Hercules, CA catalog # 1725271). The primers for *CaMK2b* (qMmuCID0021273), *CaMK2d* (qMmuCIP0030149), and *CaMK2g* (qMmuCIP0030022) were pre-validated primePCR primers from Bio-Rad. The *Camk2a* primers (AGGTGTGTGAAGGTGCTGG, and TGGAGTCGGACGATATTGGG) were designed with NCBI primer design tool (https://www.ncbi.nlm.nih.gov/tools/primer-blast/) and validated in house to recognize all splicing variants containing the kinase domain. qPCR data were analysed by the software Bio-Rad CFX Manager 3.1, using *Gapdh* expression as the loading control.

For flies, the total RNA from adults was prepared with Direct-Zol RNA miniprep kit with on-column DNase treatment (ZYMO Research, #2050). cDNA synthesis and qPCR were carried out as described above. The primers for qPCR were obtained from the FlyPrimerBank^61^ and were further validated for efficiency and specificity. Expression of RP49 was used for normalization. The primer IDs and sequences are as follows: RP49 (PD41810: AGCATACAGGCCCAAGATCG, TGTTGTCGATACCCTTGGGC), Thor (PD43730: CAGATGCCCGAGGTGTACTC, CATGAAAGCCCGCTCGTAGA), Sulfiredoxin/CG6762 (PD42226: GCATCGATGAGACCCACCTG, GATCCACAGCAGGTCGATGG), Gadd45 (PD42384: GGCCTTTTGCTACGAGAACG, CGCAGTAGTCGACTAGCTGG), Socs36E (PP11279: ATGGGTCATCACCTTAGCAAGT, TCCAGGCTGATCGTCTCTACT), GstD2 (PP27238: AAACCGCGTTTGGATTTCTCG, GTGGAGACAGTGGACAGGAT), Ank2 (PD41602: TGTGGTCATGTTAGGGTGGC, TTCAAAGCCCTTGCATTGGC), Kay (PA60087: ACTCCAACGCTTCGTACAACGATA, CACTTGAAGTATCGGTCGTGTC), Lk6 (PA60203: CAAACGCCCAGTAACATC, GCTGTAGGACCACACGCTTGAC), Atf3 (PP9314: AAGACGCCAGAGATCCTCAAC, GCAACTGGAATGACTGCTGTC), Drep3 (PP34108: GACGATGGTTTGGACGATGC, TGTTCCTCGTGATGTCCTTGA), Htk (PP1052: TACCTGGTACATTACACAGGCT, GTGCGAGTTTTCTGCTTGGA), SNF4Aγ (PP10614: ACCTCCGCCAAGTTGGTTG, CGCACACCGTTGTAGACGA), Sra (PP5475: CCGATGCACCTGATCCGAC, TTGTTCTTGCTTCTGCCGTTG), Axud1 (PP35701: GAGATAATCGTACTAGGCGATGC, GCGGAGTCAAGAATGTTGTCAA).

---

**Extended Data Fig. 1.**
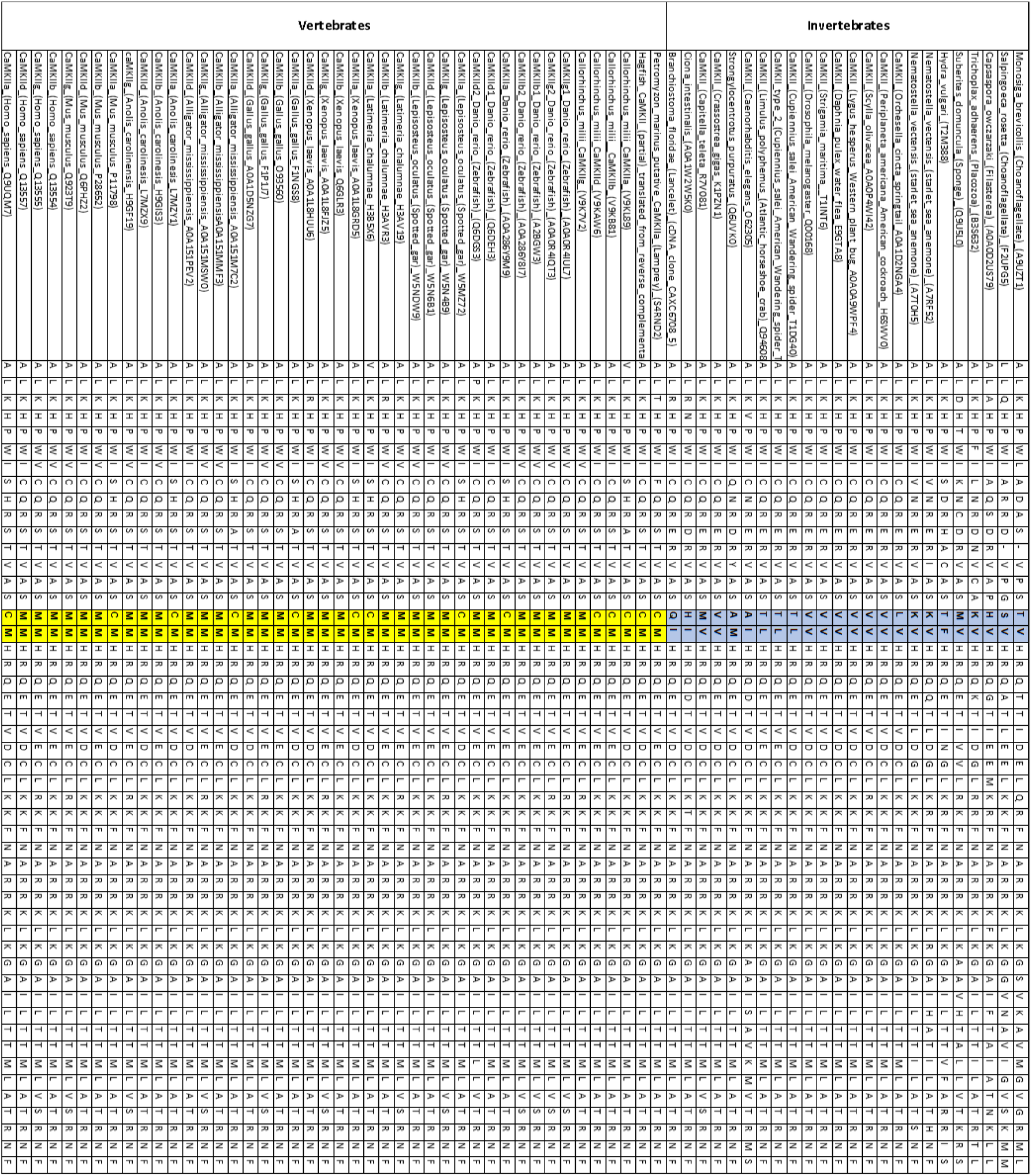
Sequence alignment of CaMKII regulatory domains reveals the emergence and conservation of ox-CaMKII in vertebrates. The paired oxidation sensitive amino acid residues (CM and MM at positions corresponding to residues 281 and 282 of mouse CaMKIIγ, highlighted yellow) are required for ox-CaMKII and are present only in the vertebrates. In contrast, none of the sampled invertebrate CaMKII possess the CM/MM pair (highlighted in light blue).

**Extended Data Fig. 2.**
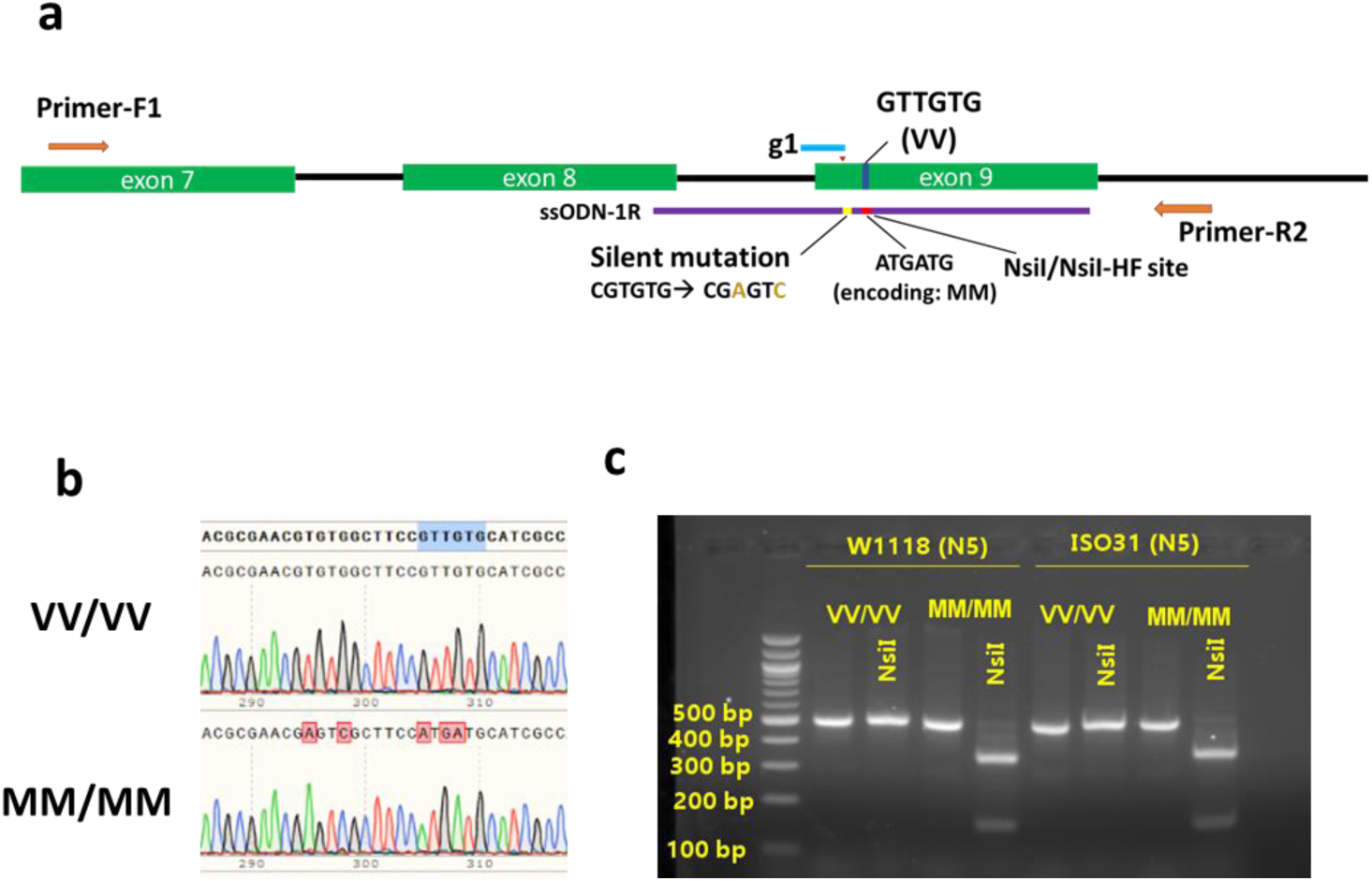
Generation of *CaMKII^MM^*/*CaMKII^MM^* flies using CRISPR. **a**, Schematics of the CRISPR guide designs and the single strand template (ssODN-1R) that mediated the homology directed recombination, resulting in mutations of codons from encoding VV to encoding MM, and introduction of the silent mutations for NsiI/NsiI-HF restriction site. Primers-F1/R2 anneal outside of the range homologous to the ssODN-1R to amplify the genomic region for genotyping. b, Chromatograms of sequencing results from the PCR products amplified by primers F1 and R2 from homozygous VV/VV (top) and MM/MM (bottom) flies. The VV/VV (wild type) and MM/MM flies are referred to as VV and MM flies in the text for brevity. c, Agarose gel electrophoresis of PCR products from flies backcrossed into the w1118 or iso31 genetic background for 5 generations. The genotypes of flies were determined by digesting the 493 bp PCR products with the restriction enzyme NsiI. PCR products amplified from the VV allele were resistant to NsiI, while those from the MM allele were cut into 345 bp and 148 bp fragments.

**Extended Data Fig. 3.**
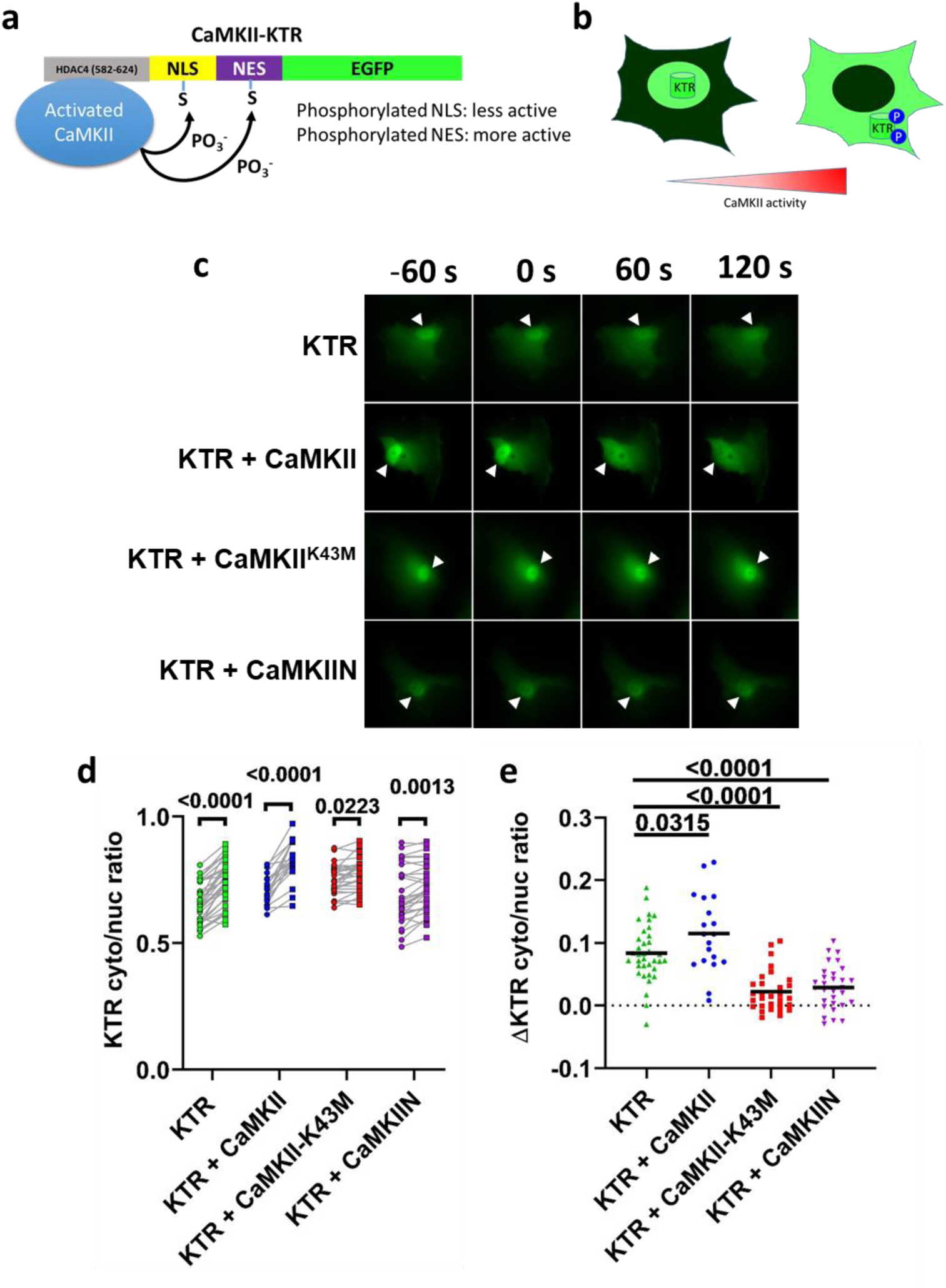
Design and validation of the CaMKII activity reporter, CaMKII-KTR. **a**, Schematic of the CaMKII kinase activity translocation reporter CaMKII-KTR (abbreviated as KTR). The N-terminus of the KTR is a well-characterized CaMKII-interacting domain from HDAC4 (AA582-624^51^), followed by a nuclear localization signal (NLS) and a nuclear exporting signal (NES). The high affinity CaMKII substrate consensus sequence (LXRXXSV) was built into both the NLS and NES (see Method). The C-terminus of the CaMKII-KTR is an enhanced green fluorescent protein (EGFP). b, The KTR shuttles between the nucleus and cytosol. Phosphorylation by CaMKII decreases the strength of the NLS while increases the strength of the NES, resulting in a net translocation of the KTR into the cytosol. The ratio between the cytosolic and nuclear signals of the KTR corresponds to the overall activity of CaMKII inside the cells. c, Fluorescent images of KTR transfected into RPE-1 cells alone, co-transfected with CaMKII, with kinase-dead CaMKII^K43M^, or with a CaMKII-specific inhibitor CaMKIIN. The arrowheads indicate nuclei. Cells were imaged at time -60 seconds (s), 0, 60 s, and 120 s, and were treated at time 0 with 50 μM of histamine. Treatment with vehicle (medium) did not elicit a response and is not shown. d, Quantification of cytosolic to nuclear KTR signal ratios in RPE-1 cells as exemplified in c immediately before and 60 seconds after the histamine treatment. *P*-values are shown in the graph, Sidak’s multiple comparisons test for repeated measurement comparing before and after histamine treatment. e, Changes in KTR cytosolic to nuclear signal ratios 60 s after histamine treatment compared to time 0 in cells shown in d. Horizontal lines indicate the means. *P*-values are shown in the graph, Dunnett’s multiple comparisons test. In (d) and (e), n = 36 KTR transfected cells, n = 19 KTR + CaMKII cells, n = 30 KTR + CaMKII^K43M^ cells, and n = 30 KTR + CaMKIIN cells. Data in (d) and (e) were from at least 2 independent experiments.

**Extended Data Fig. 4.**
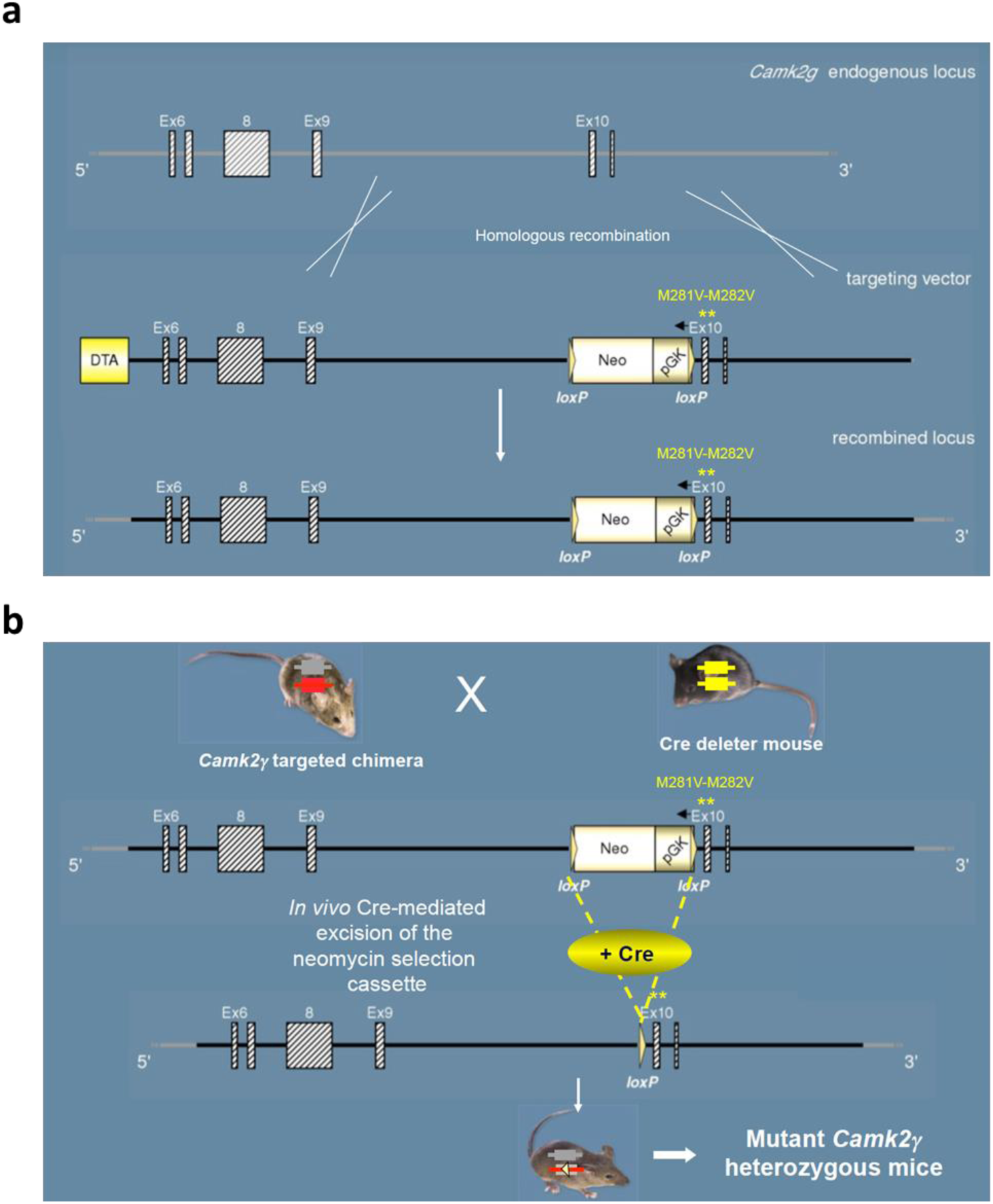
Generation of knock-in *CaMKII^VV^/CaMKII^VV^* mice. **a**, Schematics of the *CaMK2g* endogenous locus (top), targeting vector (middle), and targeted locus (bottom). The MM residues are encoded by exon 10. The gene targeting was carried out on ES cells of C57BL6/n background. b, chimeric mice bearing the targeted VV allele were crossed with Cre mice to remove the Neo-pGK cassette between the two LoxP sites. The resulting *CaMKII^MM^/CaMKII^VV^* heterozygous mice were generated backcrossed to C57BL/6J mice for >7 generations.

**Extended Data Fig. 5.**
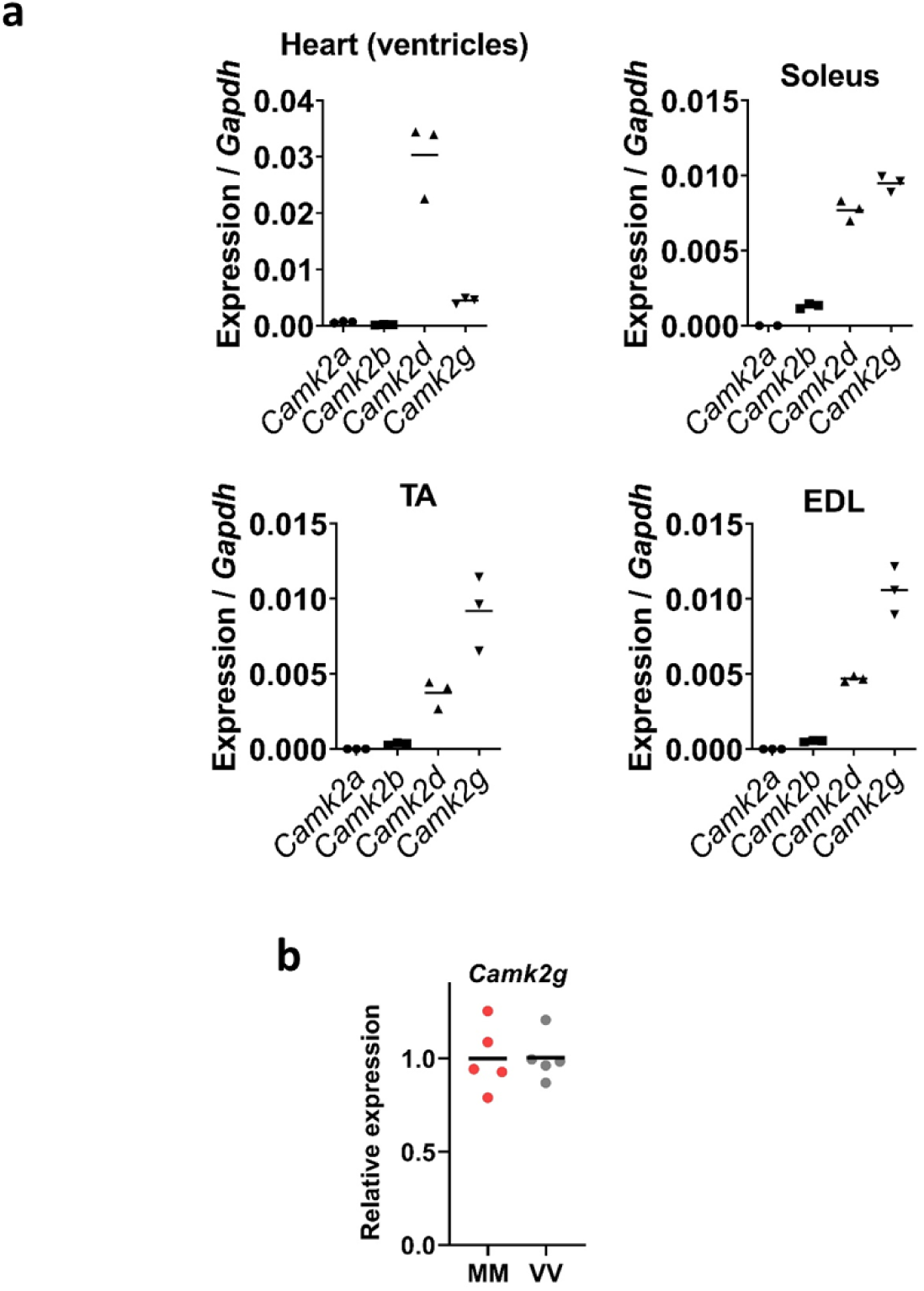
Expression of CaMKII isoforms in representative striated muscles quantified by RT-qPCR. **a**, Expression of CaMKII isoforms in the heart, soleus, TA (tibialis anterior), and EDL (extensor digitorum longus) muscles relative to *Gapdh*. Each tissue was represented by three individual wild type animals. b, RT-qPCR assay for the expression of *Camk2g* in gastrocnemius muscles from MM and VV mice; n = 5 for each genotype. In a and b, horizontal lines indicate means.

**Extended Data Fig. 6.**
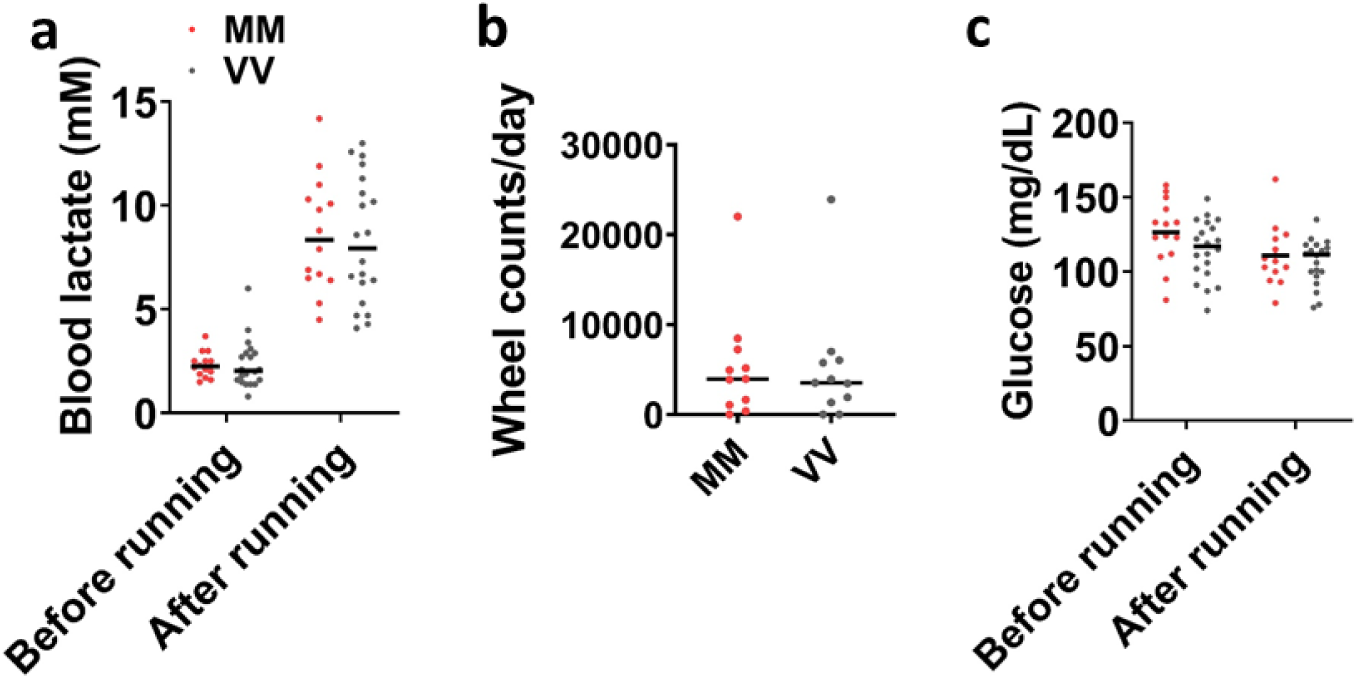
Blood lactate, glucose, and voluntary running. **a**, Lactate concentrations measured with a drop of blood from tail tips before and immediately after treadmill exercise, n = 14 MM and n = 20 VV mice, no statistically significant differences were present between genotypes either before or after exercise. b, Counts of wheel rotations during 24 hours on the 6^th^ day of running wheel access by individual MM (n = 11) and VV (n = 11) mice, no statistically significant difference was found between genotypes. c, Blood glucose concentration measured from the same mice at the same time as in (a). In a-c, horizontal lines indicate the medians.

**Extended Data Fig. 7.**
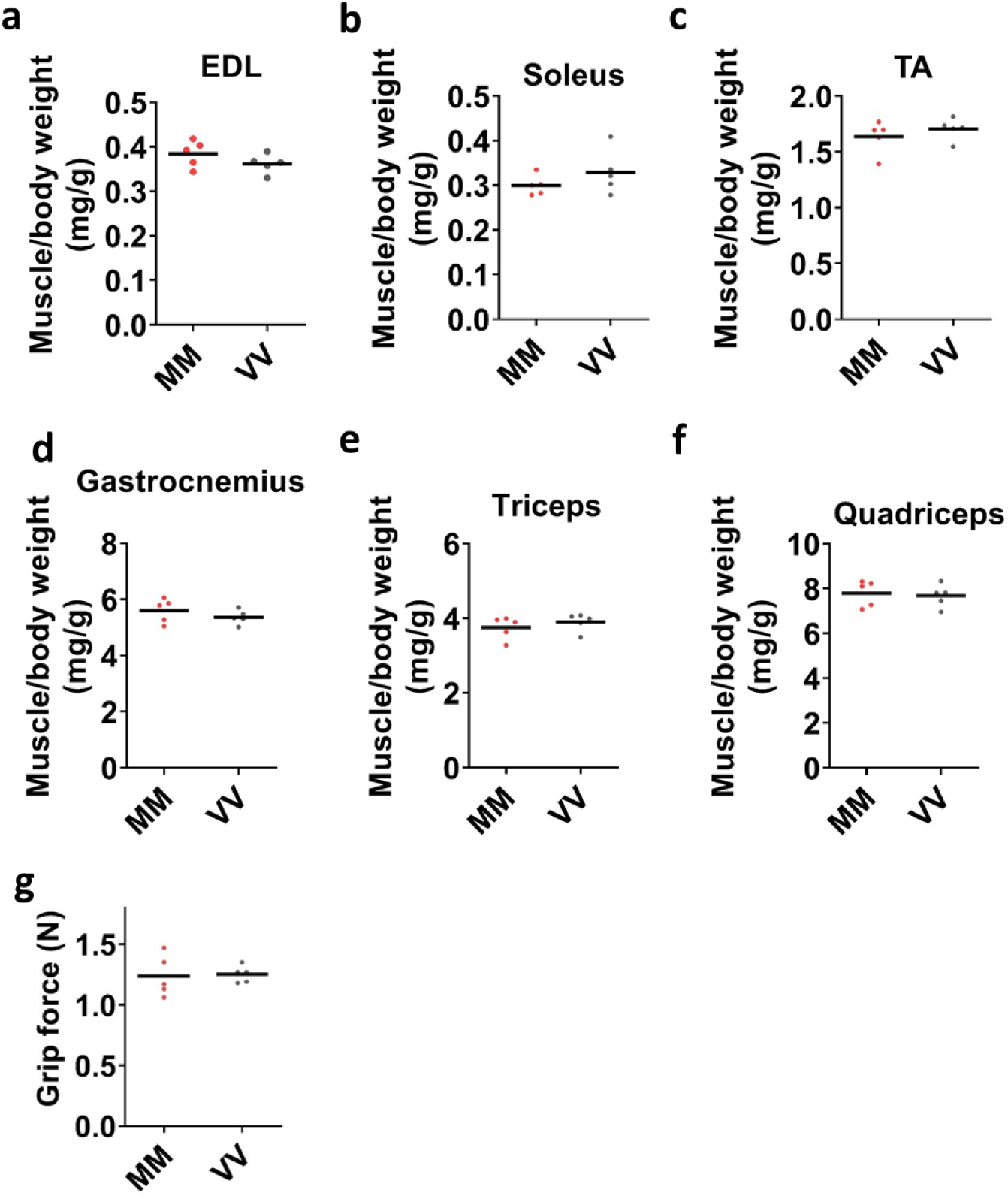
MM (WT) and VV skeletal muscles show no difference in weight and strength. **a-f**, Muscle to body weight ratios of representative skeletal muscles, a, EDL (extensor digitorum longus). b, Soleus. c, TA (tibialis anterior). d, Gastrocnemius. e, Triceps. f, Quadriceps. g, Grip force of front paws. Horizontal bars indicate means, n = 5 MM and n = 5 VV mice from (a-g).

**Extended Data Fig. 8.**
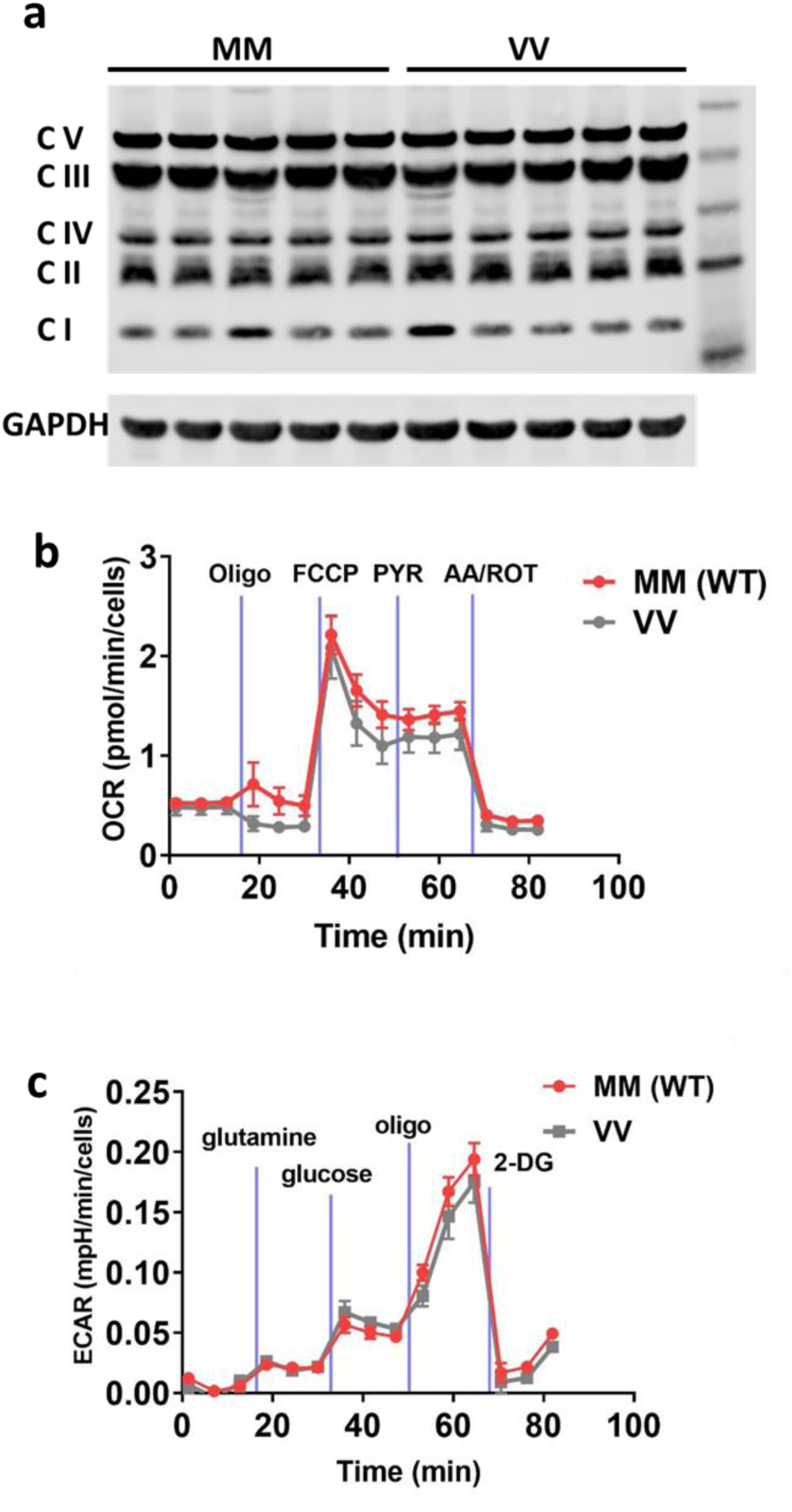
Oxidative and glycolytic capacities of MM and VV muscles are similar. **a**, Western blot of gastrocnemius muscle extracts showed no difference in the expression of representative protein subunits from mitochondrial oxidative phosphorylation complexes (n = 5 mice for each genotype). Isolated flexor digitorum brevis (FDB) fibres were analysed for (b) oxygen consumption (OCR) and (c) extracellular acidification (ECAR) rates. Points and error bars are mean ± SEM. No statistically significant (multiple t-test adjusted for false-discovery) differences were found between MM and VV fibres, n = 5 MM (WT) mice and n = 4 VV mice in b and c.

**Extended Data Fig. 9.**
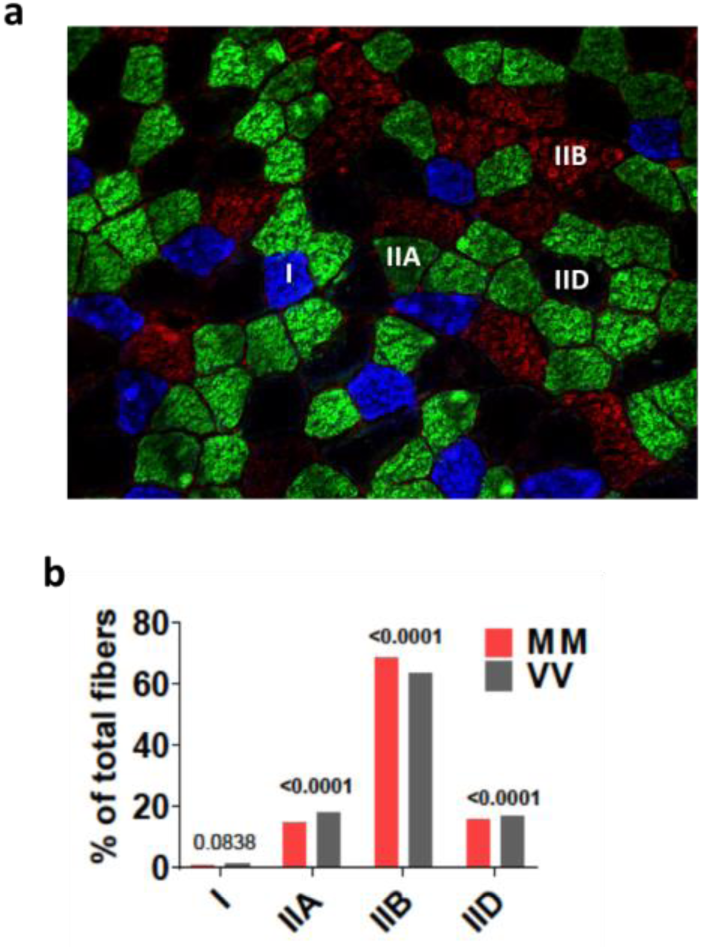
Analysis of skeletal muscle fibre types in MM and VV mice. **a**, an example cross sectional image of quadriceps muscle with immunostaining for myosin heavy chain isoforms I (blue), IIA (green), IIB (red), and IID (black, unstained). Note that this image was chosen to show the staining of all four types of myosin heavy chain isoforms, and the proportions of isoforms in this image do not reflect the entire cross section of the muscle because fibre types are not uniformly distributed throughout the cross-section of the muscles. b, quantification of the percentages of fibre types determined by myosin heavy chain isoform expression in MM (n = 5 mice, 1 section per mouse, 20110 fibres counted in total) and VV (n = 5 mice, 1 section per mouse, 22922 fibres counted in total) mice (Fisher’s exact test comparing the same fibre types between MM and VV muscles).

**Supplementary table 1.** Statistics of differential expression analysis by Cuffdiff2.

| #Categories | MM (sed) vs.<br>VV (sed) | MM (ex) vs.<br>VV (ex) | MM (sed) vs.<br>MM (ex) | VV (sed) vs.<br>VV (ex) |
| --- | --- | --- | --- | --- |
| (1) gene_exp.diff | 30529 | 30599 | 30598 | 30618 |
| (2) OK | 14628 | 14689 | 14825 | 14628 |
| (3) signif | 93 | 96 | 743 | 324 |
| (4) signif.ann | 41 | 46 | **582** | **216** |
| (5) signif.ann.fpk2 | 40 | 41 | 490 | 194 |
| (6) signif.ann.fpk2.logFC1.5 | 0 | 3 | 34 | 24 |
| (7) signif.ann.fpk5 | 33 | 31 | 365 | 157 |
| (8) signif.ann.fpk5.logFC1.5 | 0 | 3 | 26 | 21 |
| (9) signif.novel | 52 | 50 | 161 | 108 |
| (10) signif.novel.fpk2 | 29 | 27 | 77 | 44 |
| (11)<br>signif.novel.fpk2.logFC1.5 | 20 | 25 | 47 | 33 |

**Supplementary table 2.**
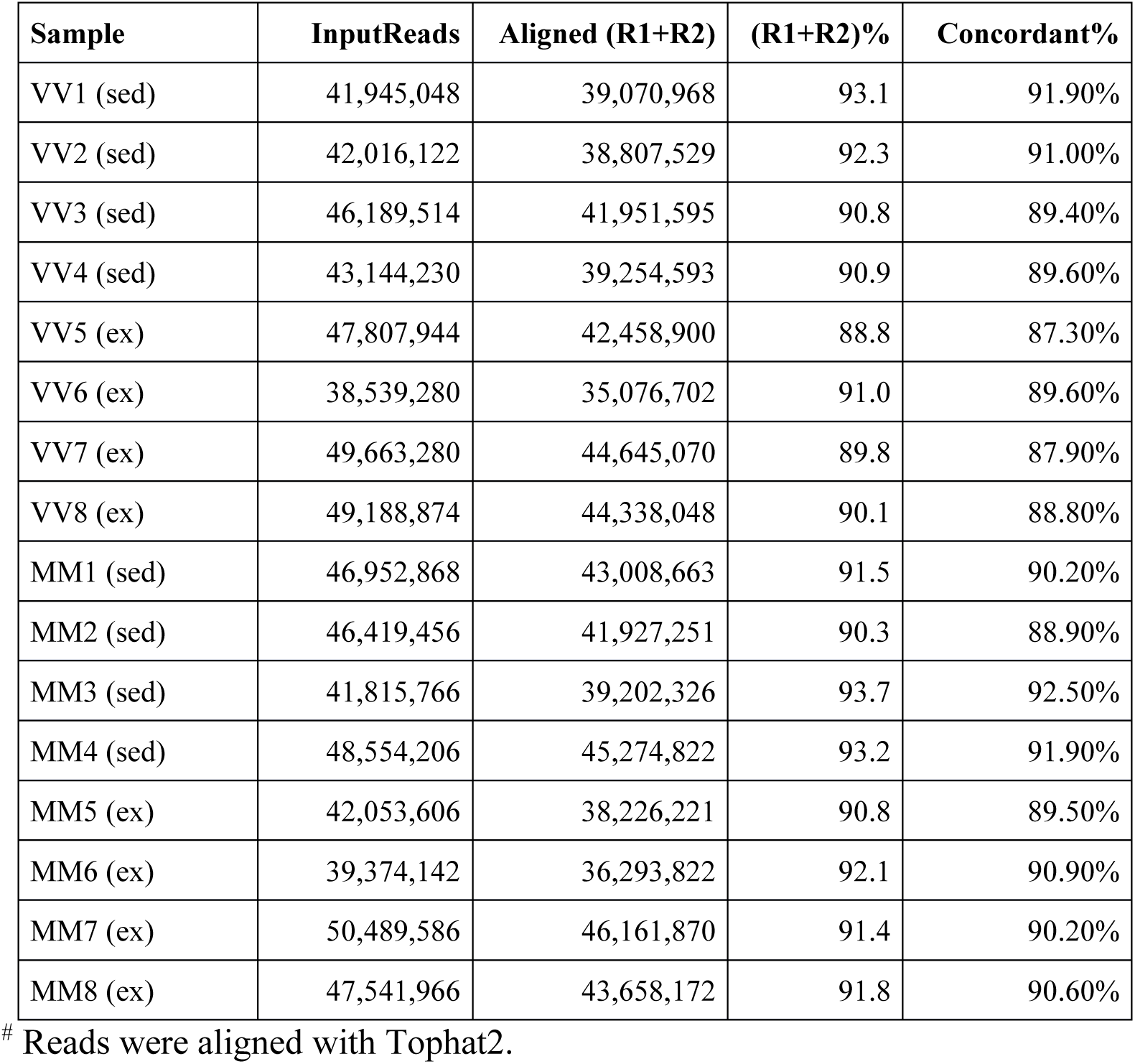
Alignment statistics of RNA sequencing reads^#^.

**Supplementary table 3.** RNA sequencing read classification.

| **Sample** | **Aligned** | **Intergenic** | **Intron** | **Exon** | **Exon-intron** | **(Ex+Ex-in)**<br>**%** |
| --- | --- | --- | --- | --- | --- | --- |
| VV1 (sed) | 39,070,968 | 10,871,032 | 1,151,817 | 20,572,924 | 6,475,195 | 69.2 |
| VV2 (sed) | 38,807,529 | 10,173,751 | 1,056,398 | 20,822,770 | 6,754,610 | 71.1 |
| VV3 (sed) | 41,951,595 | 9,850,891 | 1,100,707 | 23,295,361 | 7,704,636 | 73.9 |
| VV4 (sed) | 39,254,593 | 9,358,135 | 1,067,585 | 21,523,146 | 7,305,727 | 73.4 |
| VV5 (ex) | 42,458,900 | 9,795,942 | 1,104,903 | 23,546,149 | 8,011,906 | 74.3 |
| VV6 (ex) | 35,076,702 | 8,350,481 | 1,020,203 | 19,380,828 | 6,325,190 | 73.3 |
| VV7 (ex) | 44,645,070 | 12,762,540 | 1,450,013 | 22,758,083 | 7,674,434 | 68.2 |
| VV8 (ex) | 44,338,048 | 12,196,685 | 1,273,665 | 23,333,460 | 7,534,238 | 69.6 |
| MM1 (sed) | 43,008,663 | 14,389,489 | 1,223,644 | 20,909,218 | 6,486,312 | 63.7 |
| MM2 (sed) | 41,927,251 | 11,612,157 | 1,202,400 | 22,166,915 | 6,945,779 | 69.4 |
| MM3 (sed) | 39,202,326 | 9,690,241 | 1,091,394 | 21,601,873 | 6,818,818 | 72.5 |
| MM4 (sed) | 45,274,822 | 11,798,947 | 1,183,309 | 24,310,729 | 7,981,837 | 71.3 |
| MM5 (ex) | 38,226,221 | 9,375,367 | 1,089,510 | 20,889,204 | 6,872,140 | 72.6 |
| MM6 (ex) | 36,293,822 | 9,136,213 | 1,069,954 | 19,752,989 | 6,334,666 | 71.9 |
| MM7 (ex) | 46,161,870 | 12,957,951 | 1,406,060 | 24,129,718 | 7,668,141 | 68.9 |
| MM8 (ex) | 43,658,172 | 12,897,911 | 1,309,884 | 22,353,997 | 7,096,380 | 67.5 |

